# DualNetGO: A Dual Network Model for Protein Function Prediction via Effective Feature Selection

**DOI:** 10.1101/2023.11.29.569192

**Authors:** Zhuoyang Chen, Qiong Luo

## Abstract

**Motivation:** Protein-protein Interaction (PPI) networks are crucial for automatically annotating protein functions. As multiple PPI networks exist for the same set of proteins that capture properties from different aspects, it is a challenging task to effectively utilize these heterogeneous networks. Recently, several deep learning models have combined PPI networks from all evidence, or concatenated all graph embeddings for protein function prediction. However, the lack of a judicious selection procedure prevents the effective harness of information from different PPI networks, as these networks vary in densities, structures, and noise levels. Consequently, combining protein features indiscriminately could increase the noise level, leading to decreased model performance.

**Results:** We develop DualNetGO, a dual network model comprised of a classifier and a selector, to predict protein functions by effectively selecting features from different sources including graph embeddings of PPI networks, protein domain and subcellular location information. Evaluation of DualNetGO on human and mouse datasets in comparison with other network-based models show at least 4.5%, 6.2% and 14.2% improvement on Fmax in BP, MF and CC Gene Ontology categories respectively for human, and 3.3%, 10.6% and 7.7% improvement on Fmax for mouse. We demonstrate the generalization capability of our model by training and testing on the CAFA3 data, and show its versatility by incorporating Esm2 embeddings. We further show that our model is insensitive to the choice of graph embedding method and is time- and memory-saving. These results demonstrate that combining a subset of features including PPI networks and protein attributes selected by our model is more effective in utilizing PPI network information than only using one kind of or concatenating graph embeddings from all kinds of PPI networks.

**Availability and implementation:** The source code of DualNetGO and some of the experiment data are available at: https://github.com/georgedashen/DualNetGO.

**Supplementary Information:** Supplementary data are available at *Bioinformatics* online.

## 1 Introduction

Proteins are the main players in biological processes, and their functions can be categorized into three aspects by Gene Ontology (GO): biological process (BP), molecular function (MF) and cellular component (CC) (Aleksander *et al*., 2023). Knowing a protein’s function helps explain its role and evaluate its importance in a biological process, and is also useful for enzyme and drug design (Radivojac *et al*., 2013). However, by 2023 less than 1% of over 200 million known proteins have been revealed their functions (Uniprot, 2023) because experimental protein annotation is laborious, time consuming and costly (Luck *et al*., 2020). Thus, automatically annotating protein functions becomes a meaningful and yet challenging task.

With the development of the Critical Assessment of Functional Annotation (CAFA) community, dozens of advanced algorithms have been proposed for automatic protein function annotation (Zhou *et al*., 2019). Some algorithms utilize sequence features that learned from neural networks (Kulmanov *et al*., 2018; Kulmanov and Hoehndorf, 2021; Cao and Shen, 2021) or protein language models (Wang *et al*., 2023; Oliveira *et al*., 2023), or from predicted structural information (Gligorijević *et al*., 2021; Boadu *et al*., 2023). In comparison, network-based methods utilize protein-protein interaction (PPI) networks to predict protein functions. PPI networks provide additional information into how proteins work cooperatively to exert a certain function, which is difficult to determine directly from protein sequences or structures.

According to the STRING database (Szklarczyk *et al*., 2023) there are seven types of evidence to define an interaction between two proteins: *neighborhood, fusion, cooccurence, coexpression, experimental, database* and *textmining*. Most of existing network-based methods use all types of PPI networks to compute a weighted summing network (Mostafavi *et al*., 2008) or an integrated graph embedding vector for each protein (Cho *et al*., 2016; Gligorijević *et al*., 2018), or use a combined PPI network that integrates edges from all evidence (Fan *et al*., 2020; Wu *et al*., 2023). As different networks vary in density and connectivity, simply combining all networks into a single one can lead to information loss (Cho *et al*., 2016). Indiscriminate use of these networks can further increase the noise level of the data and result in decreased model performance (Bi *et al*., 2023), especially when some of the included networks or features are less relevant than others to the downstream task (Mostafavi *et al*., 2008). How to properly and effectively utilize different PPI networks is still to be explored for protein function prediction.

Recently, Maurya et.al. proposed a feature selection strategy to handle heterogenous graph data, where features of neighbors at different hops may not correlate with node features, which hampers the performance of classical graph neural network (GNN) models on node classification tasks (Maurya *et al*., 2023). Their proposed method intelligently determined a suitable combination of features derived from the same graph. Furthermore, this strategy can be applied to a boarder range of problems beyond the GNN models. The problem of utilizing information from different PPI networks is a good example of such an extension.

To better utilize different PPI networks, we develop a dual network model named DualNetGO, extended from the existing feature selection strategy, to predict protein function by effectively determining the combination of features from PPI networks and protein attributes without enumerating each possibility. We design a feature matrix space that includes eight matrices: seven for graph embeddings of PPI networks from different evidence and one for protein domain and subcellular location. After encoding each PPI network into low-dimensional latent factors, the two multilayer perceptron (MLP) components of DualNetGO, the Classifier and the Selector, are trained alternately to evaluate the importance of each matrix and choose a suitable combination to predict protein functions. Experiment results show that DualNetGO outperforms other network-based methods on the human and mouse datasets and is insensitive to the choice of graph embedding methods. Further evaluation shows that with proper settings DualNetGO takes less time and requires less memory in data preprocessing and training. These results demonstrate that DualNetGO is an efficient and effective network-based model for protein function prediction by using different PPI networks, providing insight into better ways of utilizing heterogeneous PPI network data.

The contributions of our work include:

1. DualNetGO achieves SOTA performance on protein function prediction over other single species PPI network-based methods. It also makes the best prediction on the CC aspect on the CAFA3 test set among all methods under comparison.
2. To the best of our knowledge, our work is the first attempt to investigate a suitable combination of graph features of PPI networks from different types of evidence for a single species, and demonstrate the effects of choosing different PPI networks on protein function prediction for different GO aspects.
3. We have conducted a comprehensive study to evaluate the effects of different graph embedding methods on various PPI networks for protein function prediction.
4. Our feature selection strategy can be applied to general scenarios where multi-modal features exist and each feature is represented as a matrix, not only for network information.

## 2 Materials and methods

### 2.1 Dataset

PPI data are retrieved from STRING database on STRINGv11.5 for human and mouse. For a specific evidence if there is a positive score, the interaction between two proteins is considered to exist. PPI networks are transformed into weighted adjacency matrices and minmax-normalized. GO functional annotations are downloaded from the gene ontology website (version 2022-01-13 release). Protein attributes that include subcellular location and the Pfam protein domain annotation are retrieved from the Uniprot database (v3.5.175), and those having fewer than 6 proteins are removed. Following the CAFA challenge setting (Jiang *et al*., 2016), we only retain protein functions with experimental evidence ‘IDA’, ‘IPI’, ‘EXP’, ‘IGI’, ‘IMP’, ‘IEP’, ‘IC’ or ‘TA’ as one-hot encoded labels, and define proteins with annotations before 2018-01-01 as the training set, those between 2018-01-02 and 2020-12-31 as the validation set and those after 2021-01-01 as the test set. This temporal holdout method to split data was proposed in the CAFA challenge to mimic a real-life application scenario instead of random splitting (Jiang *et al*., 2016). To make sure there are a sufficient number of proteins for each label, we retain labels with at least 10, 5 and 1 proteins in the training, validation and test set, respectively, following a previous study (Wu *et al*., 2023). To further reduce the correlation or dependency between GO terms, any labels containing more than 5% of the number of proteins in human and mouse PPI network are removed as well. The statistics of the final training, validation and test set, and different PPI networks, are shown in Supplementary Table S1,S2.

Furthermore, to compare with other state-of-the-art methods such as NetGO3.0 (Wang *et al*., 2023) and DeepGOplus (Kulmanov and Hoehndorf, 2021) which are not PPI network-based models, we downloaded the CAFA3 dataset from the TEMPROT paper (Oliveira *et al*., 2023) for large-scale multi-species training and testing. More details about data collection and preprocessing can be found in Supplementary Section 15.

### 2.2 Method

#### 2.2.1 Transformer-based autoencoder for PPI and protein attributes

DualNetGO contains two components: a graph encoder and a predictor (Figure 1a,b). The graph encoder is a previously published transformer-based autoencoder (denoted as **TransformerAE**) (Wu *et al*., 2023) that takes protein attributes and PPI networks as input and outputs low-dimensional embeddings. We choose TransformerAE for its superior performance in integrating networks and features without the message-passing mechanism of graph neural networks to better capture complex network properties. In the TransformerAE, the adjacency matrix and protein attribute matrix together go through 6 multi-head attention layers for the encoder and another 6 layers for the decoder to fuse information from the two sources. The core of the attention mechanism is the Scaled Dot-Product Attention (Vaswani *et al*., 2017), where *Q* is query, *K* is key and *V* is value matrix, and *d*_*k*_ is the dimension of query and key vectors in the matrix.

**Fig 1:**
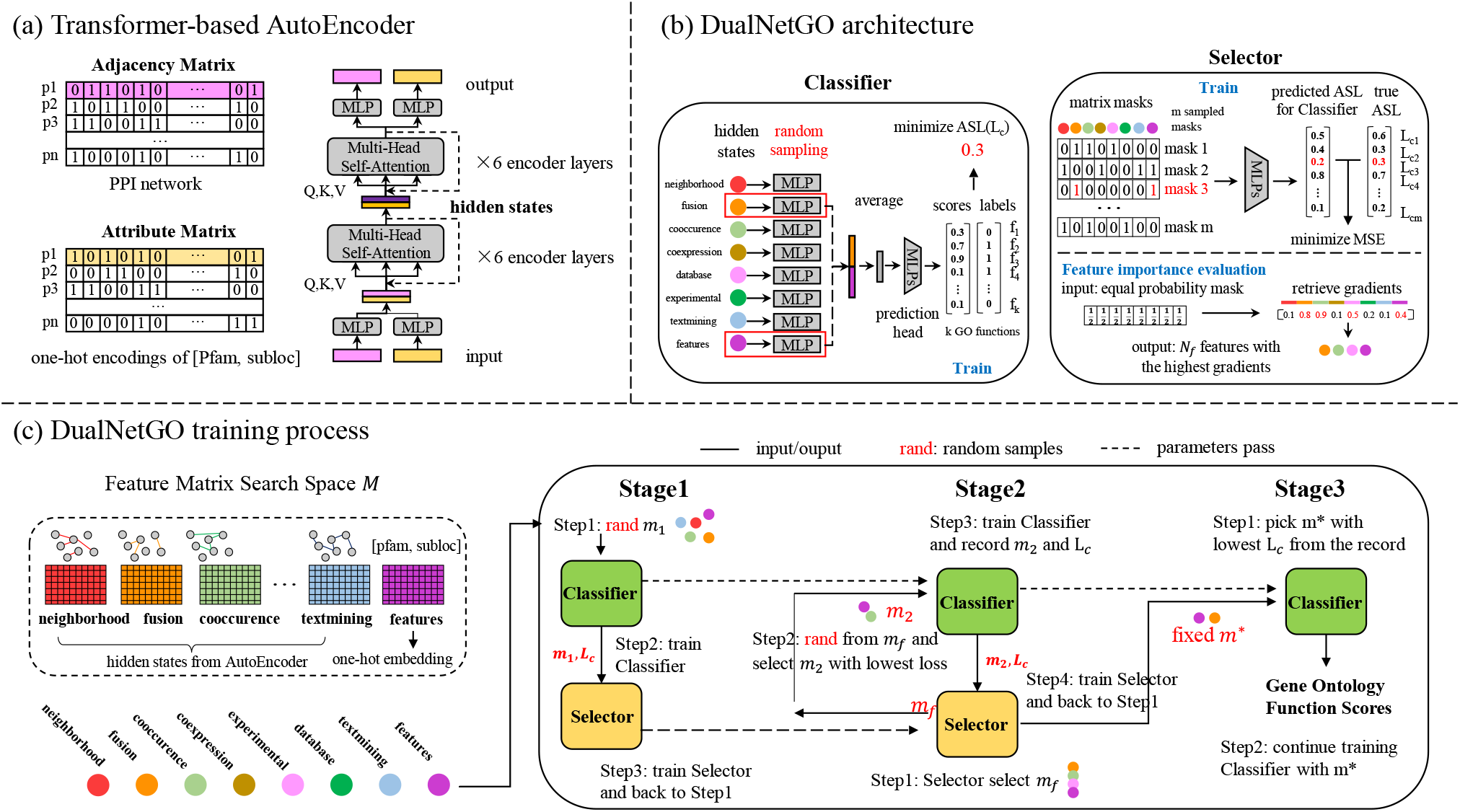
The workflow of DualNetGO. (a) Architecture of TransformerAE for generating PPI network graph embeddings. (b) Architecture of the Classifier and the Selector component of DualNetGO. In the Classifier section, the randomly sampled features (eg. *fusion* and *features*) are indicated by red blocks, and only them pass through the following MLP modules. In the Selector section, the selected features are indicated by values of 1 in red colors in a matrix mask. The corresponding predicted ASL from the Selector and the true ASL from the Classifier are also in red color. The values below the colored ribbon representing the absolute values of gradients of the trained Selector with respect to elements in the mask input. The highest values are in red color and the associated features will be selected for futher sampling. (c) Training process of DualNetGO to select features for protein function prediction. ASL: asymmetric loss. MSE: mean squared error.

Scaled Dot-Product Attention:

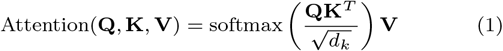

During the self-supervised learning process of TransformerAE, differences between the original input before being passed into the encoder and the reconstructed output after the decoder are minimized by binary cross-entropy. Only the hidden states for PPI networks are included in the feature matrix space.

#### 2.2.2 Dual network architecture of DualNetGO

The predictor is a dual network comprised of a *classifier* and a *selector* (Figure 1c).

##### Classifier

The Classifier takes features as input and outputs scores for each GO function. It maintains a one-layer MLP with ReLU as the nonlinear activation function for each matrix in the feature matrix space to further reduce the dimension, and a two-layer MLP (denoted as the **prediction head**) with softmax activation function in the second layer to output a score for each GO term. Selected feature matrices will first go through their own MLP modules and be averaged, then pass through the prediction head.

In the Classifier, asymmetric loss (ASL) (Ridnik *et al*., 2021) is used as the loss function to reduce the contribution of easy negative samples, encouraging the model to make more positive predictions in a multi-label task with imbalanced samples across classes.

ASL is defined as:

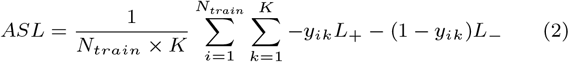

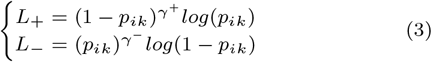

*N*_*train*_ is the number of proteins in the training set, K is the number of functions in a specific category, *γ*^+^ and *γ*^−^ are the focusing parameters for positive and negative samples, respectively. When both parameters are set to 0, ASL is equivalent to a binary cross entropy. In this study, we set *γ*^+^ to 0 and *γ*^−^ to 2, the same as the default setting in the ASL paper and those used in *CFAGO*. Results with different *γ*^−^ can be found in Supplementary Table S3.

##### Selector

The Selector is a two-layer MLP for selecting a set of important feature matrices based on the gradients of the model, to further narrow down possible feature combinations. The input is a one-hot encoded feature mask representing the selected feature matrices for input to the Classifier. For example, if the *coexpression* and *textmining* networks are selected, the mask is [0,0,0,1,0,0,1,0], where a value 1 indicates the selection of the corresponding feature matrix. The output is a scale value approximating the validation loss of the Classifier. Previous work suggests that the absolute values of gradients of a trained machine learning model can be used to evaluate the importance of the corresponding element in the input (Hechtlinger, 2016). By training with various masks and their corresponding validation losses from the Classifier, the Selector serves as a surrogate function to the Classifier. The vector input to the Selector supports the evaluation of feature importance by gradients. Specifically, the Selector learns to evaluate the Classifier’s performance with the selected subset of feature matrices, and is expected to output a lower value when the selected input subset is more suitable for predicting protein function.

Mean squared error (MSE) is used as the loss function between the predicted loss and the true loss from the Classifier on the validation set.

#### 2.2.3 Training process of DualNetGO

The training process is divided into three stages, and two networks are trained alternately in the first two stages. The number of training epochs for each stage is defined as *E*1, *E*2 and *E*3, respectively, and the maximum number of feature matrices to be included for prediction is defined as *N*_*f*_. These numbers are set as hyperparameters before training.

##### Stage 1

A random combination of feature matrices is sampled from the matrix space *M* with no more than *N*_*f*_ matrices at the beginning of each epoch, with a mask as *m*_1_. These matrices go through the Classifier, and an ASL prediction loss for protein functions on validation set *L*_*c*_ is calculated and backpropagated. The mask *m*_1_ and the loss *L*_*c*_ is used as input for the Selector, and an MSE loss *L*_*s*_ is calculated between the output of the Selector and *L*_*c*_, and backpropagated. After *E*1 epochs the Selector learns to evaluate Classifier’s performance with a given mask. This stage can be viewed as an **exploration** process to gather information to train the Selector as a good surrogate function to the Classifier, which requires a variety of mask vectors and their corresponding validation losses from the Classifier.

##### Stage 2

In each epoch we first create a mask with equal weights of 0.5 representing an equal chance for each matrix to be selected, and then use this mask as input for the Selector and calculate the gradient of each element in the mask. Given a trained Selector model, the absolute values of gradients of the input are expected to reflect the importance of each element. We select the corresponding matrices with top *N*_*f*_ absolute gradient values, indicated by the indices in the mask, to form a new matrix space *M*_*f*_. Because the optimal combination could be a subset of *M*_*f*_, a fixed number of combinations are further sampled from *M*_*f*_ and evaluated by the Classifier on the validation set to narrow down the range of optimal combinations. The lowest validation loss and the corresponding mask *m*_2_ are recorded, and *m*2 is used for a similar process in Stage 1 to train the Classifier and then the Selector. This stage can be viewed as an **exploitation** process, which utilizes the information from Stage 1 by retrieving the gradients in the Selector to determine a set of features with the highest importance.

##### Stage 3

The mask with the minimal validation loss across the record is identified as *m*^*^, and the training process for the Classifier is continued by only using the corresponding matrices with *m*^*^. Only weights in the MLP modules with respect to *m*^*^ and the prediction head in the Classifier will be updated for *E*3 epochs. Performance on test set data is reported as the final results when the Classifier achieves the best Fmax score on the validation set.

### 2.3 Evaluation metrics

We use three protein-centric metrics (Macro-F1 (M-F1), F1, accuracy, Fmax) and two term-centric metrics including two types of area under the precision-recall curve (AUPR), micro-AUPR (m-AUPR) and macro-AUPR (M-AUPR), to evaluate prediction performance. Accuracy is defined as the proportion of proteins with all functions correctly predicted. The threshold is determined by achieving the highest Fmax score on the validation set. m-AUPR is the AUPR calculated across all true labels and predictions, and M-AUPR is the average of AUPR values over AUPRs of all GO terms.

Fmax is the official metric of CAFA competition and defined as:

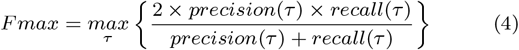

where *τ* is a flexible threshold for both recall and precision to obtain the maximum Fmax score.

Precision and recall for a multi-label task are defined as:

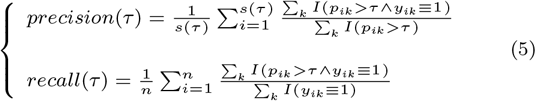

*s*(*τ*) denotes the number of proteins that are predicted with at least one function. *k* is the total number of labels for a specific functional category. *p*_*ik*_ is the predicted score for the function and *y*_*ik*_ is the ground truth with 1 indicating the existence of the function. *n* is the total number of proteins to be evaluated.

### 2.4 Experiment setup

The architecture of DualNetGO and the training procedure including learning rates and weight decays are based on previous work (Maurya *et al*., 2023) or by manual search (Supplementary Table S4,S5). Overall, as the Classifier and the Selector are trained alternately in each epoch of Stage 1 and 2, the Classifier is trained for *E*1 + *E*2 + *E*3 epochs, and the Selector is trained for *E*1 + *E*2 epochs. We set *E*2 + *E*3 = 100 for ease of implementation with an early stopping strategy and limit *N*_*f*_ ranging from 2 to 5. Different from the previous work using k-hop features as the feature matrix space to deal with heterophilic graph data, we include hidden states from seven PPI networks as potential features to address the multi-network utilization problem. Without losing generality of our model, we also include a protein attribute matrix that has a different number of dimensions from the other matrices to handle potentially multi-modal data.

We compare DualNetGO with two baseline methods and four network-based models.

**Naive**. The naive method simply assigns the relative frequency of a term over all proteins in the training set as the score for this term for all proteins in the test set.

**BLAST**. The BLAST method transfers GO terms of a target protein in the training set to the query protein in the test set via the *blastp* software, and the identity score of alignment is used as a coefficient for all assigned terms.

**Mashup**. This is a linear and shallow model that uses a matrix factorization-based approach to compute low-dimensional vectors for proteins across diffusion states from different PPI networks (Cho *et al*., 2016). Mashup cannot extract the complex and nonlinear information in various PPI networks.

**deepNF**. This is a deep learning model to construct a compact low-dimensional representation from complex topological properties of PPI networks. It first uses a separate MLP module for the PPMI matrix of each network to reduce the dimensionality, and then concatenates these low-dimensional features. To fuse information from different networks, it utilizes an MLP-based autoencoder (MLPAE) structure to construct an integrated low-dimensional representation from concatenated features (Gligorijević *et al*., 2018).

**Graph2GO**. This model utilizes variational graph autoencoders (GAE) on a combined PPI network and a sequence similarity network, with protein attributes as input features (Fan *et al*., 2020).

**CFAGO**. This method designs a transformer-based autoencoder (denoted as TransformerAE) to cross-fuse the combined PPI graph and protein attributes with the attention mechanism(Wu *et al*., 2023).

All hyperparameters of the TransformerAE graph encoder and data preprocessing follow the *CFAGO* paper (Wu *et al*., 2023). All experiments are conducted with a single RTX 3090 GPU with 24G memory. Details can be found in Supplementary Section 3 and 6.

## 3 Experiments

### 3.1 DualNetGO outperforms competing network-based models

Figure 2 shows that DualNetGo outperforms other models on most of the metrics across GO aspects and organisms, except for MF in mouse. Specifically, DualNetGO gains improvement of at least 0.045, 0.062 and 0.142 (up to 0.459, 0.226 and 0.464) in terms of Fmax for BP, MF and CC respectively on human, and 0.027, 0.077 (up to 0.296 and 0.502) for BP and CC on mouse. For m-AUPR, DualNetGO achieves at least 0.058, 0.026 and 0.141 higher for BP, MF and CC respectively on human, and 0.001, 0.147 for BP and CC on mouse. Improvements in M-AUPR, M-F1 and F1 can also be observed in half of the scenarios (more details in Supplementary Table S6,S7).

**Fig 2:**
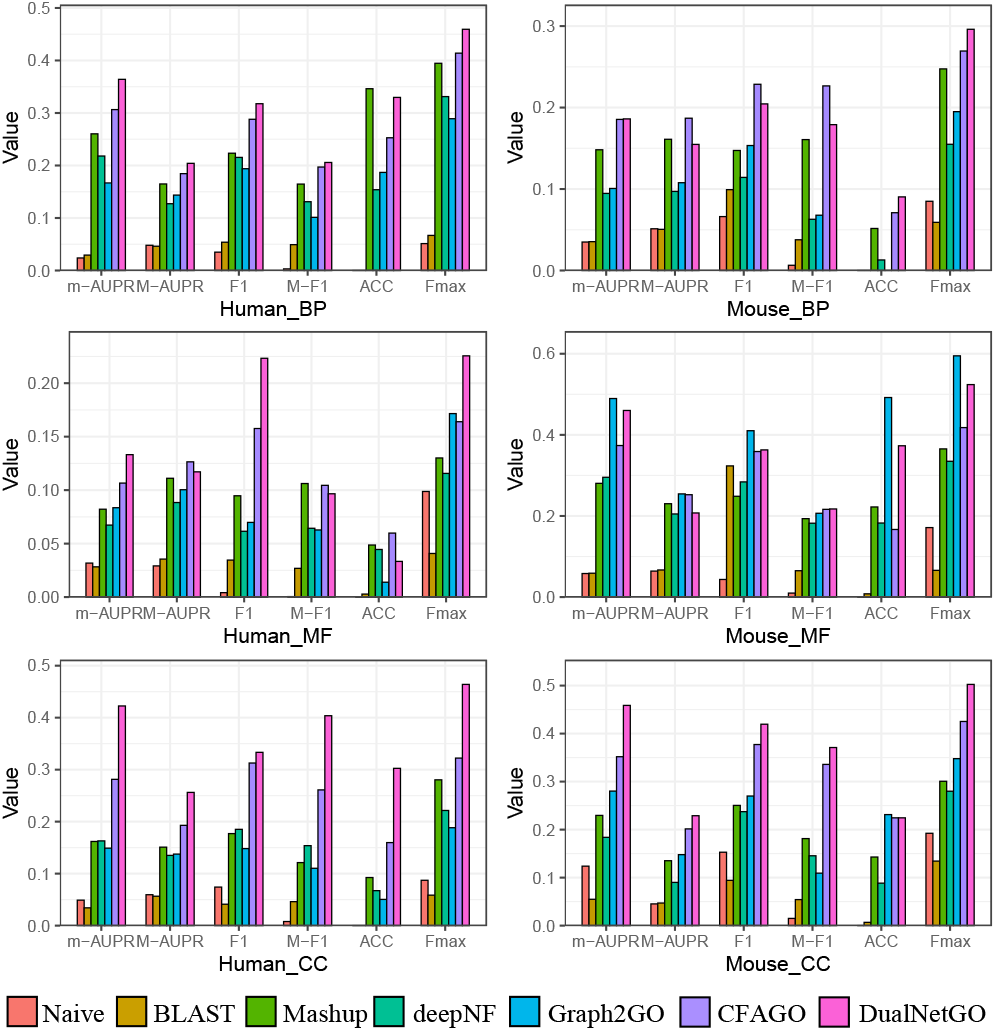
Performance of DualNetGO for protein function prediction compared with other network-based models.

In the MF category of mouse, DualNetGO produces slightly worse results than Graph2GO, a model that also utilizes sequence similarity network which is not included in other models, in addition to the PPI network. Several studies (Fan *et al*., 2020; Oliveira *et al*., 2023) suggest that MF is more related to sequence patterns that may not be reflected by Pfam protein domains and PPI networks.

The accuracy of DualNetGO is not distinct compared to other metrics. A reason could be that the threshold for accuracy is determined by the evaluation on the validation set with the highest Fmax score, but the data distribution between the validation set and the test set may be different.

These observations of DualNetGO demonstrate that feature selection across different PPI networks is an effective strategy to improve protein function prediction performance.

### 3.2 DualNetGO benefits from other graph embedding methods

We implement other graph embedding methods including node2vec (Grover and Leskovec, 2016), GAE (Kipf and Welling, 2016) and MLPAE to replace the TransformerAE in the PPI network preprocessing step, in order to investigate whether DualNetGO is affected by the choice of graph embedding methods. To make a more comprehensive comparison, we also consider situations when only using protein attributes without PPI networks (denoted as **Feature** in figures), using Esm2 sequence embeddings (**Esm2**) (Lin *et al*., 2023), using randomly generated latent factors (**random**), and not using any graph embedding techniques but only the raw adjacency matrix (**NoEmbed**) as input for each PPI network. The results of DualNetGO are the same as previously reported by selecting features from the seven PPI networks and the UniProt protein attributes. Figure 3 shows that the effects of graph embedding methods differ by PPI usage scenarios, but DualNetGO outperforms other scenarios on the human dataset. The superior performance of DualNetGO is not affected by the choice of graph embedding methods, as we can see that DualNetGO performs better than the best scenario of other methods, regardless of the choice of graph embedding methods in all three categories, except for the use of node2vec in BP. Similar observation is found on the mouse dataset (Supplementary Figure S2). These results show the effectiveness of DualNetGO on protein function prediction with the matrix selection strategy is robust and could benefit from advanced graph embedding algorithms in the future.

**Fig 3:**
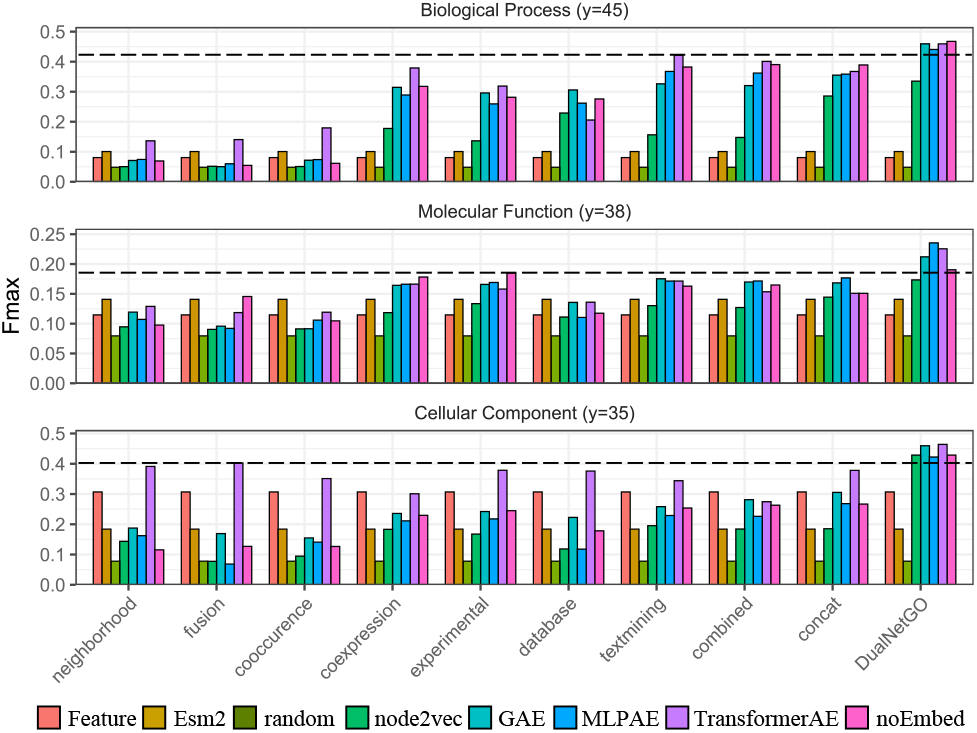
Fmax scores across different graph embedding methods and different PPI usage settings on human dataset. Second best Fmax are indicated by dash lines.

### 3.3 The contribution of the Selector and different training stages

To demonstrate the importance of each component of DualNetGO and the design of a three-stage training procedure, we conduct ablation experiments on the Selector and both Stage 1 and Stage 2 of the training process individually. More details about designs of different ablation tests can be found in Supplementary Section 10.

Table 1 shows that all components contribute to the superior performance of DualNetGO. Stage 1 is the most important in performance, reflected by dramatic Fmax drops in most scenarios. The Stage 1 is important because the Selector need to be trained first with the validation loss provided by the Classifier in Stage 1 to produce accurate evaluation of the importance of each feature matrix in Stage 2. Since the evaluation is based on the gradients of the Selector model, without Stage 1 the Selector will be randomly initiated, and thus the gradients will be irrelevant to the feature importance. While in Stage 2 the Selector may gradually produce accurate evaluation as more and more combinations are sampled for training, this alternative is not as efficient as that in Stage 1. The reason is that the combination sampling in Stage 2 depends on the Selector, which is not fully random. Therefore, only a limited combinations will be sampled in Stage 2 with the same number of epoch as Stage 1.

**Table 1.**
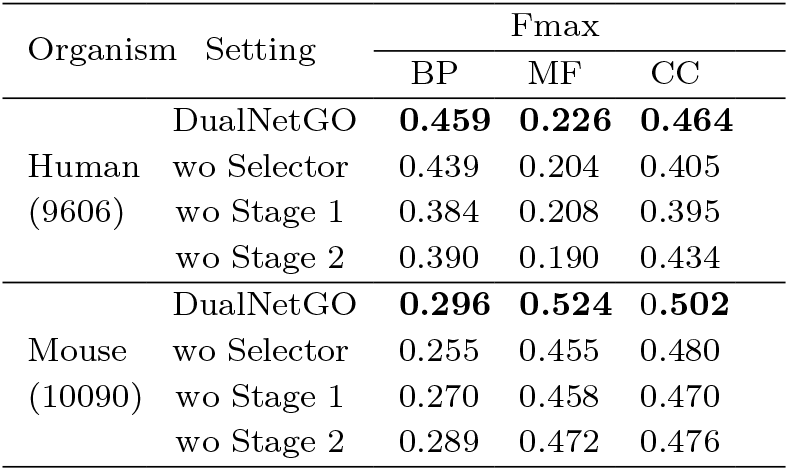
Ablation test on each component of DualNetGO.

Both the Selector and Stage 2 (representing an exploitation process) play a role in the effectiveness of DualNetGO. Without the Selector, there is no guidance for the Classifier to choose a suitable subset of the features for prediction. Using the mask with respect to the lowest validation loss of the Classifier in the stochastic training of Stage 1 results in reduced performance, as the optimal combination may not be sampled. Even the optimal subset is sampled, with a few epochs of training it is less likely to produces the lowest validation loss, thus will not be selected for fixed training in Stage 3. We also demonstrate that training the Classifier with all features without the Selector results in much worse performance than with the Selector (Supplementary Section 11). In Stage 2, a fixed number of combinations are further sampled with the subset selected by the Selector, and only the optimal combination with the lowest validation loss inferred by the Classifier will be used to further train the Classifier and the Selector. Because in Stage 2 the features are selected based on the gradients of the Selector, and there are additional inference rounds to select the best mask, the training process is less random and less stochastic than Stage 1, which can be viewed as a refinement process.

### 3.4 Analysis of training time and validation loss

The update of parameters in the Classifier can be viewed as a stochastic process, which strikes a balance between determining the optimal subset and reducing the training time. While fluctuating loss is observed during stage 1 and 2 (Figure 4), the overall training loss decreases over time and Fmax on the validation set is reaching the plateau.

**Fig 4:**
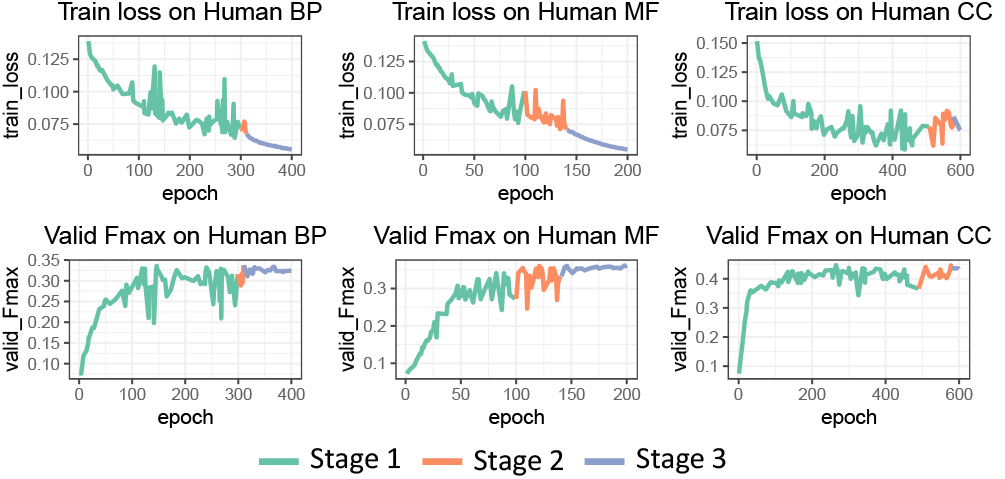
Training loss and validation Fmax of DualNetGO across different training stages on the human dataset.

Time for preprocessing data and training is also compared for various models. Results (Figure 5) show that the data preprocessing time needed for DualNetGO TransformerAE and CFAGO is more than one magnitude longer than the other models. This is because the two models adopt the same graph embedding method TransformerAE, which is much larger (up to 82 million) than other models (Supplementary Table S9). The long running time for TransformerAE is due to the choice of a large number of epochs in the original paper, but in practice a much smaller number of epochs such as 500 is applicable. Furthermore, TransformerAE is not the only option for DualNetGO, with GAE and node2vec among the most time-efficient graph embedding algorithms. Graph2GO, which performs well on mouse MF, is the second time consuming model in preprocessing PPI data, due to the exhaustive sequence alignments between any two sequences out of about 20,000 sequences and the choice of using 100 epochs to train the GAE.

**Fig 5:**
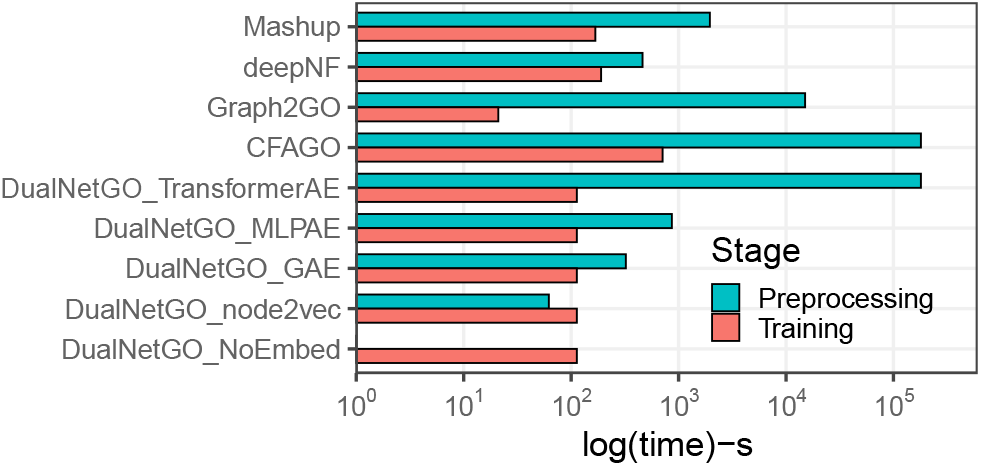
Comparison of preprocessing and training time across models.

As DualNetGO adopts a heuristic strategy to determine the combination instead of enumerating each possibility, a careful choice of the hyperparameters of the epochs in stages 1 and 2 may be necessary to approach the optimal solution, as suggested by another dual-network model that deals with heterophilic graph data (Maurya *et al*., 2022). Fortunately, the performances across hyperparameters show an obvious pattern (Supplementary Section 12), and the search of hyperparameters costs little time. Overall, DualNetGO is still more efficient in determining a suitable combination of features than enumerating all possibilities.

### 3.5 Embedding-centric setting versus evidence-centric setting for DualNetGO

We denote the previous search space for searching across different PPI networks given a specific graph embedding method (TransformerAE) as the **embedding-centric** setting. We call another way of utilizing DualNetGO model the **evidence-centric** setting, which searches across different graph embedding algorithms given a PPI network from a specific type of evidence (Figure 6). As an extension to the DualNetGO, we also implement the model in the evidence-centric setting using only the combined PPI network and evaluate the performance on both human and mouse dataset. Collectively, DualNetGO with embedding-centric and evidence-centric settings produces the top-2 results in almost all metrics in all datasets except in mouse MF where Graph2GO generally performs better (Supplementary Table S6). We also evaluate the use of other graph embedding methods for the embedding-centric model and other PPI networks for the evidence-centric model, and find even better results. The best performance and the corresponding parameters can be found in Supplementary Table S10,S11.

**Fig 6:**
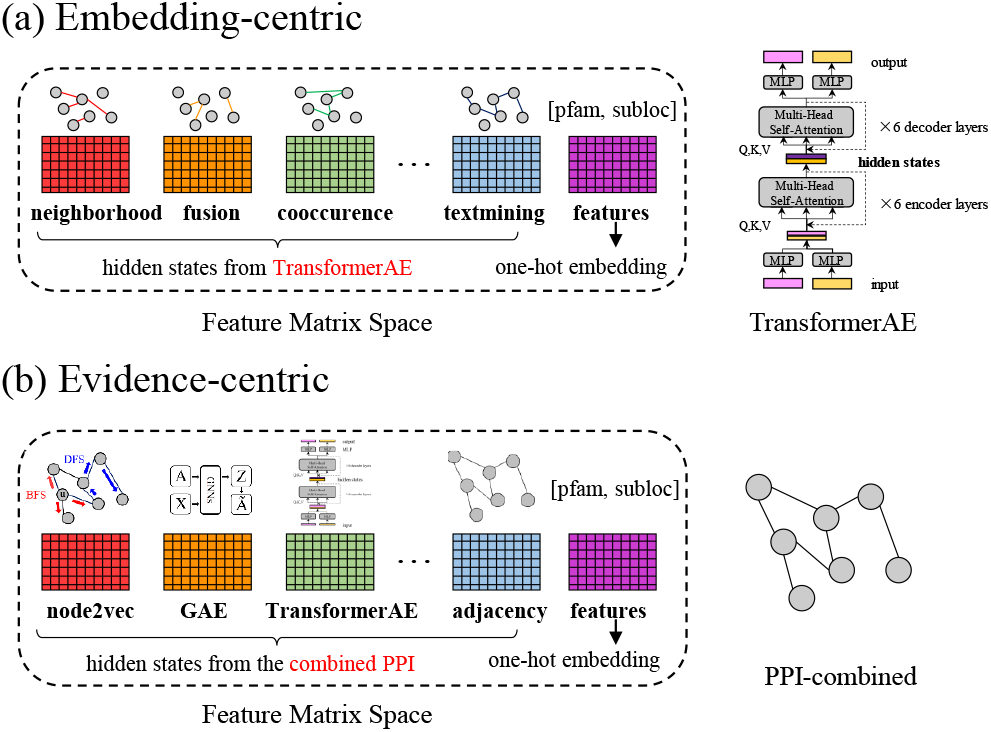
Comparison of preprocessing and training time across models.

### 3.6 Comparison with other protein function prediction models on CAFA3 test set

To compare DualNetGO with other state-of-the-art methods on the CAFA3 test set and demonstrate its generalization capability, we train our model on the CAFA3 training set under a multi-species setting. To show the versatility of our model to integrate multi-modal features, we incorporate sequence embeddings encoded by a protein language model Esm2 (Lin *et al*., 2023) in the feature selection space instead of the original Pfam/subloc features, while the TransformerAE is still trained by fusing the network adjacency matrix and the Pfam/subloc attribute matrix. In addition, the homology search strategy, which ensembles DualNetGO and BLASTp predictions, can be also applied. We denote models without and with homology search as DualNetGO and DualNetGO+, respectively. More details can be found in Supplementary Section 15-17.

For comparison we choose models that harness information from different sources. DeepGOCNN (Kulmanov *et al*., 2018), TALE (Cao and Shen, 2021) and TEMPROT (Oliveira *et al*., 2023) are sequence-based methods, TransFun (Boadu *et al*., 2023) is a structure-based model, DeepGraphGO (You *et al*., 2021) is a multi-species network-based model, NetGO3.0 (Wang *et al*., 2023) and DeepGOplus (Kulmanov and Hoehndorf, 2021) are ensemble models. Results for the Naive, BLASTp, DeepGOCNN, TALE and TEMPROT are directly cited from the TEMPROT paper as they are also evaluated on the CAFA3 test set. For TransFun we use the predicted scores provided by its authors and evaluate the performance under our CAFA3 settings. For DeepGraphGO we train the model using the provided script and training set. For NetGO3.0 we use the online server. For DeepGOplus we use the provided model weights.

Results (Table 2) show that DualNetGO and DualNetGO+ produce the higest Fmax and AUPR scores on CC, comparable results on BP, and worse results on the MF aspect. Similar results are also observed on the previous filtered human/mouse datasets using the DualNetGO model trained on CAFA3 data (Supplementary Section 18). DualNetGO’s superior performance on CC suggests that protein functions on CC are more related to PPI networks than sequences, whereas functions on MF largely depend on sequence properties. This conjecture is also supported by the observation that the Esm2 sequence embedding feature is not selected by DualNetGO as one of the final features for CC, but instead the textmining and cooccurence networks are selected. The Esm2 embedding features are selected for both BP and MF (Supplementary Section 19). Especially, the homology search strategy plays a more important role for the improvement of MF than those of BP and CC. The correlations between PPI network and CC, and that between sequences and MF are also supported by another study (Ibtehaz *et al*., 2023).

**Table 2.**
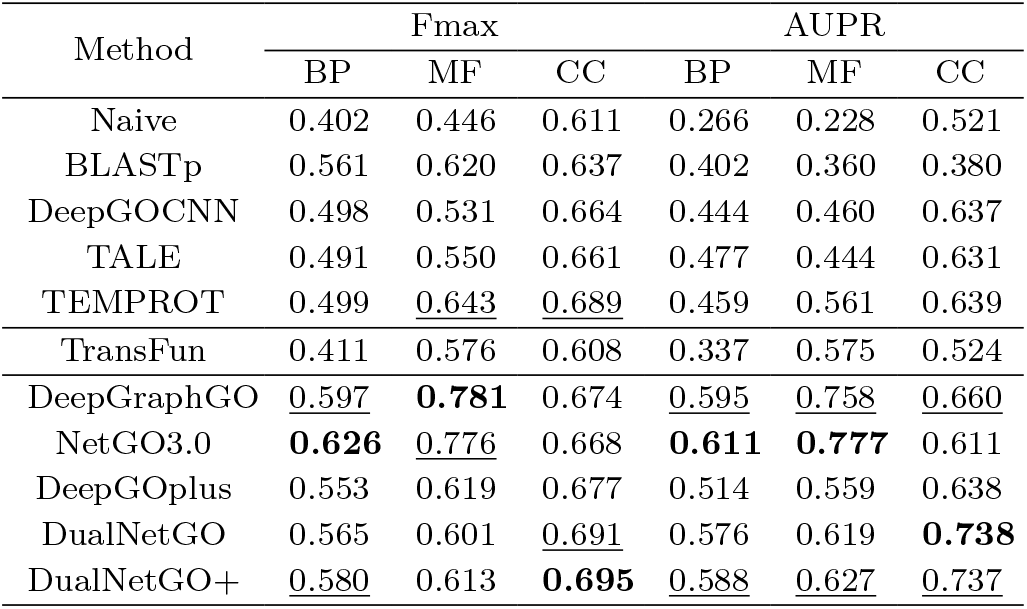
Evaluation of different methods on the CAFA3 multi-species test set. Highest results are in bold, with second and third highest underlined.

| Method | Fmax |  |  | AUPR |  |  |
| --- | --- | --- | --- | --- | --- | --- |
|  | BP | MF | CC | BP | MF | CC |
| Naive | 0.402 | 0.446 | 0.611 | 0.266 | 0.228 | 0.521 |
| BLASTp | 0.561 | 0.620 | 0.637 | 0.402 | 0.360 | 0.380 |
| DeepGOCNN | 0.498 | 0.531 | 0.664 | 0.444 | 0.460 | 0.637 |
| TALE | 0.491 | 0.550 | 0.661 | 0.477 | 0.444 | 0.631 |
| TEMPROT | 0.499 | <u>0.643</u> | <u>0.689</u> | 0.459 | 0.561 | 0.639 |
| TransFun | 0.411 | 0.576 | 0.608 | 0.337 | 0.575 | 0.524 |
| DeepGraphGO | <u>0.597</u> | **0.781** | 0.674 | <u>0.595</u> | <u>0.758</u> | <u>0.660</u> |
| NetGO3.0 | **0.626** | <u>0.776</u> | 0.668 | **0.611** | **0.777** | 0.611 |
| DeepGOplus | 0.553 | 0.619 | 0.677 | 0.514 | 0.559 | 0.638 |
| DualNetGO | 0.565 | 0.601 | <u>0.691</u> | 0.576 | 0.619 | **0.738** |
| DualNetGO+ | <u>0.580</u> | 0.613 | **0.695** | <u>0.588</u> | <u>0.627</u> | <u>0.737</u> |

### 3.7 Analysis of features selected for predicting protein functions

As DualNetGO can determine a suitable subset of PPI for protein function prediction, the chosen combination reflects the importance of different PPI networks. For the embedding-centric model, we implement five different graph embedding algorithms and count the occurrences of each PPI network in these experiments. As shown in Figure 7a, textmining provides the most valuable information in BP prediction for both human and mouse, and CC is more related to protein attributes (denoted as **feature**). CC’s close relation to protein attributes is not surprising because the protein attributes used in this study include Pfam domain and subcellular location, and subcellular location could cause information leakage due to its close relation to CC. In the evidence-centric model, the number of graph embeddings chosen as features is counted across different PPI networks. In Figure 7b we observe that TransformerAE generally performs better than the other methods, demonstrating its superior performance. However, the simpler deep learning autoencoder, MLPAE, is also effective on extracting PPI information for MF prediction. We also perform a thorough hyperparameter search on E1, E2 and num feat select on all three aspects, and retrieve the features selected by DualNetGO. We then count the frequency of each feature matrix being selected by DualNetGO models with top-10 Fmax scores for BP, MF and CC aspect. Results in Figure 7c show that For BP the best result is produced by combining the hidden states from the textmining network and Esm embeddings. For MF the best result is produced by combining the hidden states from the database network and Esm embeddings. For CC the best result is produced by combining the hidden states from cooccurrence and textmining networks. This phenomenon also suggest that CC is more related to the PPI network property than the sequence property.

**Fig 7:**
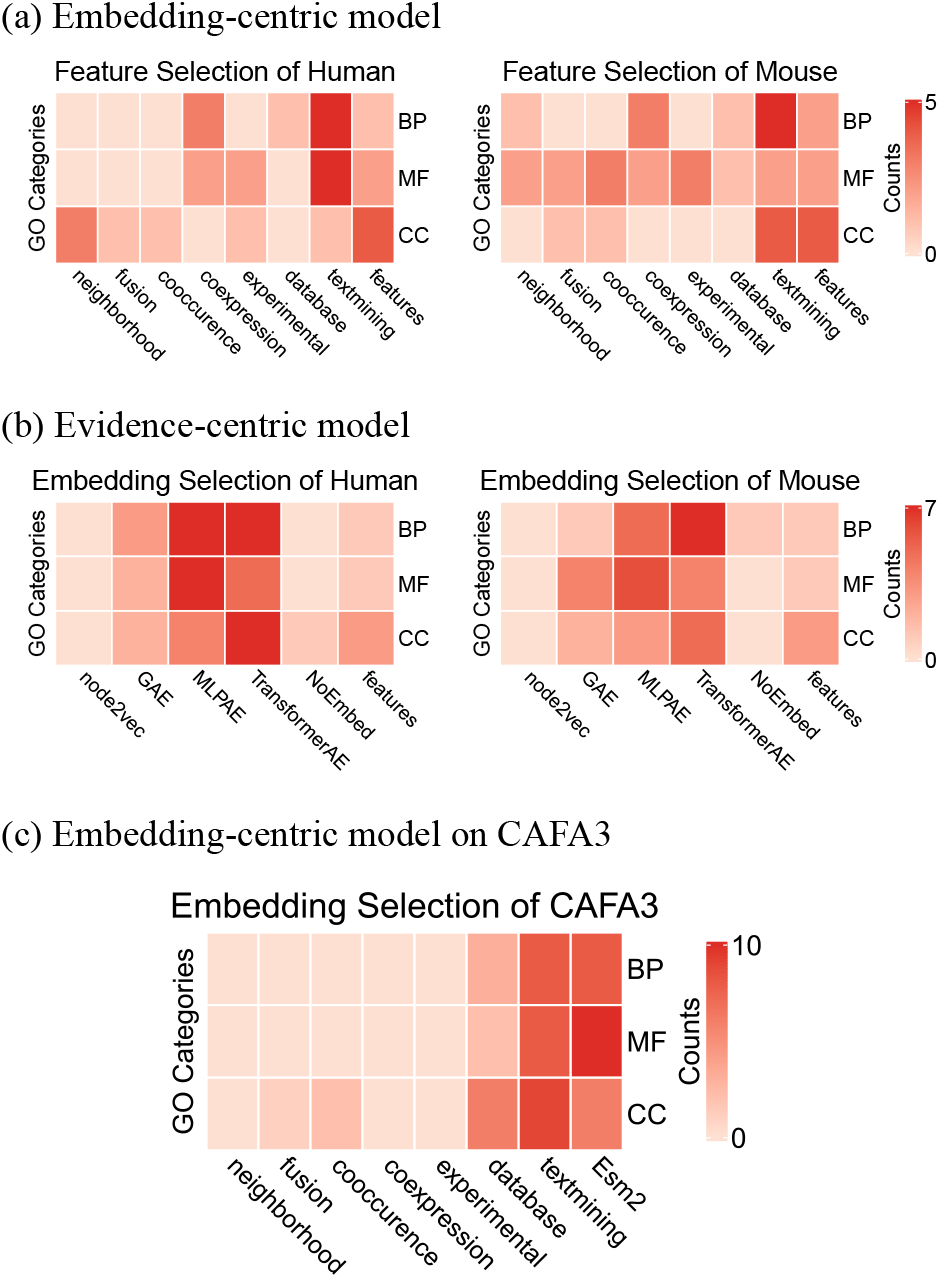
Feature importance across different settings of DualNetGO.

### 3.8 Effect of the “selection during training” strategy

The DualNetGO model is trained by alternate evaluation of feature selection and update of classifier’s weights. Before the best feature combination is determined, all features contribute to the classification loss during the feature sampling process. Therefore, even though a certain feature matrix is not included in the final features and not used for prediction, it helps update the classifier’s weights in stage 1 and stage 2, and attributes to the improvement of the model. This gives DualNetGO the advantage of fully utilizing all collected data for training the classifier but only using an optimal subset of them for prediction. To demonstrate the effectiveness of the “selection during training” strategy, we enumerate all possible combinations of feature matrices and train a corresponding classifier using each of the combinations. Results in Table 3 show that the best results from all feature combinations (denoted as **Enumerate**) are worse than those generated from DualNetGO (denoted as **Select**) in the filtered human/mouse datasets. Similar results are found in the CAFA3 multi-species dataset except a slightly higher AUPR socre on the MF aspect for the Enumerate model, shown in Table 4.

**Table 3.**
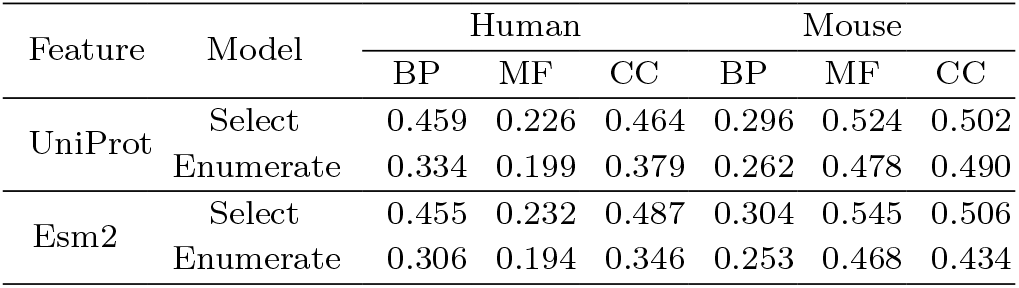
Comparison between Fmax scores from DualNetGO feature selection training and from the best results out of all feature combinations on the filtered datasets.

**Table 4.**
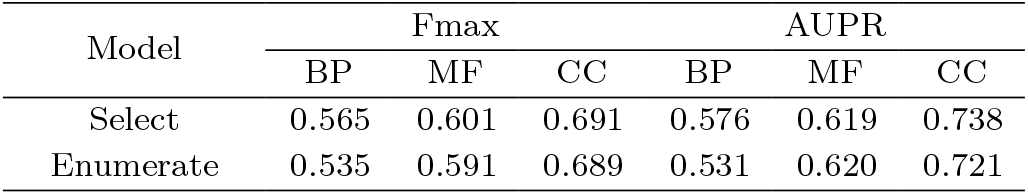
Comparison between performance from DualNetGO feature selection training and from the best results of all feature combinations on the CAFA3 test set.

We also find that the optimal combination of features selected by the “selection during training” strategy of DualNetGO is not the same as the set of features that are found by enumeration to provide the best results. Most of them are overlapped, indicating that the DualNetGO model plays the feature selection role (Table S18, 19). Comparing the features selected by DualNetGO and by Enumerate when using Esm2 embeddings as protein attributes instead of the original Pfam+subloc information retrieved from UniProt, we find that the same set of features are selected for the CC aspect on the human dataset. However, the performance of DualNetGO is better than that of Enumerate, which directly demonstrates the importance of incorporating other features in the training process for better performance. In addition, we find that using Esm2 embeddings instead produces higher results than using UniProt annotations by DualNetGO except on the BP aspect of human, but Esm2 is only selected as the final feature for human CC and mouse MF. Furthermore, the Esm2 embeddings themselves may not be as predictive as the original UniProt annotations, as the best results from enumerating combinations of Esm2 embeddings are worse than those from enumerating combinations of UniProt annotations. These phenomena suggest that features of lower relevance can also be utilized by DualNetGO to improve its performance.

## 4 Discussion

The results of this study demonstrate that DualNetGO outperforms other single-species, PPI network-based methods on protein function prediction on all aspects, and makes better predictions on the CC aspect for the CAFA3 test set. Our model’s intelligent matrix selection strategy takes full advantage of all training data to improve the performance, even if some features are not selected as the final features and not used for prediction. Our experiments show that DualNetGO yields better results than any combinations of matrices in the feature selection space. In addition, DualNetGO’s superior performance is insensitive to the choice of graph embedding methods, which makes it a versatile framework for dealing with multi-modal data when additional information of proteins such as embeddings from protein language models, knowledge graphs, and 3D structures are available. Our comprehensive study, which evaluates the effects of different graph embedding methods on different PPI networks for protein function prediction, provides valuable insight for future research. Furthermore, as a model with feature selection mechanisms, DualNetGO indicates that CC is more related to PPI networks, and MF depends more on sequence properties. However, how the performance of graph embedding methods on protein function prediction is related to the properties of different PPI networks, which PPI evidence to pay more attention to are both opening questions for future exploration.

One limitation of DualNetGO is that it does not support end-to-end training at the current stage, which means the overall performance would largely depend on the qualities of all features in the feature selection space. Previously we found that the good performance of DualNetGO was insensitive to graph embedding methods for human and mouse. However, for less common species that lack sufficient PPI or Pfam information, the graph hidden states generated by the self-supervised TransformerAE model may not provide sufficient information for protein function prediction. This limitation is reflected by the worse performance on the BP and MF aspects than another multi-species model DeepGraphGO. Also, other features that exploit sequence properties, such as those used in NetGO3.0 and DeepGOplus, must be included in the feature selection space to improve the MF performance. In the meanwhile, the non-end-to-end training procedure of DualNetGO creates versatility to effectively combine different information sources and better facilitates for multi-species training and prediction, which is generally lacked by other PPI network-based models. Another drawback is that the training set of network-based models is usually smaller than those used by other models, because only proteins recorded in the PPI network are retained. As a result, some representative proteins may not be fully utilized to train the model. This issue will be alleviated as more and more PPI data is collected.

For further improvement of the model, more advanced graph embedding methods to preprocess PPI networks and more sophisticated network structures than MLPs for prediction can be adopted. One can try to train the graph encoders and DualNetGO by an end-to-end manner, or include various high quality features from other studies in the feature selection space.

