## Supplementary for "DualNetGO: A Dual Network Model for Protein Function Prediction via Effective Feature Selection": Supplementary.docx

**Supplementary materials**

Here we provide more details about the experiment designs and results in this paper.

**1. Statistics of different datasets**

After filtering out frequent, dependent GO terms and those with few proteins, we obtain the human and mouse dataset with relatively small numbers of terms but each with a sufficient number of proteins.

| Organism | Statistics | BP | | | | | MF | | | | | CC | | | | |
| --- | --- | --- | --- | --- | --- | --- | --- | --- | --- | --- | --- | --- | --- | --- | --- | --- |
|  |  | train | | valid | | test | train | | valid | | test | train | | valid | | test |
| Human  (9606) | # protein | 3197 | | 304 | | 182 | 2747 | | 503 | | 719 | 5263 | | 577 | | 119 |
|  | # GO terms | 45 | | | | | 38 | | | | | 35 | | | | |
| Mouse  (10090) | # protein | 2714 | | 336 | | 155 | 1185 | | 232 | | 126 | 4014 | | 694 | | 147 |
|  | # GO terms | 42 | | | | | 17 | | | | | 37 | | | | |
| CAFA3  Multi-species | # protein | 47691 | 5252 | | 2392 | | 32421 | 3587 | | 1137 | | 45309 | 4985 | | 1265 | |
|  | # GO terms | 3992 | | | | | 677 | | | | | 551 | | | | |

**Table S1.** Statistics of proteins and labels in different datasets. Taxonomy codes are included in parentheses. The first two large rows are for the filtered human/mouse dataset with data downloaded from the 2022 UniProt dataset. The last large row is for the CAFA3 dataset with filtered GO terms.

**2. Statistics of different PPI networks**

Here we provide some statistics on different PPI networks to give an overview of their heterogeneity on topology.

| PPI Networks | Human (9606) | | | | | Mouse (10090) | | | | |
| --- | --- | --- | --- | --- | --- | --- | --- | --- | --- | --- |
|  | Nodes | Edges | Density | Degree | Clustering | Nodes | Edges | Density | Degree | Clustering |
| neighborhood | 19385 | 410,412 | 2.18e-3 | 21.17 | 0.06 | 21373 | 491,904 | 2.17e-3 | 23.08 | 0.05 |
| fusion |  | 22,898 | 1.22e-4 | 1.18 | 0.01 |  | 22,876 | 1.01e-4 | 1.07 | 0.01 |
| cooccurrence |  | 61,142 | 3.25e-4 | 3.15 | 0.10 |  | 76,214 | 3.35e-4 | 3.58 | 0.10 |
| coexpression |  | 6,525,628 | 0.035 | 336.63 | 0.25 |  | 7,794,954 | 0.034 | 365.67 | 0.29 |
| experimental |  | 4,950,896 | 0.026 | 255.40 | 0.20 |  | 7,827,412 | 0.034 | 367.19 | 0.25 |
| database |  | 401,438 | 2.14e-3 | 20.71 | 0.35 |  | 3,820,662 | 0.017 | 179.23 | 0.36 |
| textmining |  | 10,169,440 | 0.054 | 524.60 | 0.18 |  | 10,882,386 | 0.048 | 510.50 | 0.21 |
| combined |  | 11,938,498 | 0.063 | 615.86 | 0.20 |  | 14,496,358 | 0.064 | 680.04 | 0.26 |

**Table S2.** Statistics of different PPI networks.

Definitions of some metrics for PPI network:

$$\text{A}\text{verage }\text{C}\text{lustering }\text{C}\text{oefficient}=\frac{1}{N}\sum_{i}^{N} \frac{L}{degree(i)(degree(i)-1)} \text{(L:\#edegs, N:\#nodes)}$$

$$\text{Average Degree}=\frac{\sum_{i}^{N} degree(i)}{N} \text{(N:\#nodes)}$$

$$\text{Density}=\frac{L}{N(N-1)/2} \text{(L:\#edegs, N:\#nodes)}$$

**3. Implementations of graph embedding methods**

Codes for implementing and evaluating this model can be found at <https://github.com/georgedashen/DualNetGO>.

For Mashup, we use the code <http://cb.csail.mit.edu/cb/mashup> and modify it to accommodate our input. The dimension of the final vectors is set to 512.

For deepNF we implement a Pytorch version based on the description and hyperparameters provided by the original paper. We use all seven types of evidence for PPI networks, whereas the original deepNF paper uses six except *textmining*.

For Graph2GO we also implement a Pytorch version but reduce the training epoch for GAE from 200 to 100. *NeighborSampler* is used for batch training to reduce input size.

For CFAGO we completely use the original code and data, including the large number of 5000 training epochs for self-supervised training.

**4. Effects of different gamma_neg values of ASL loss on protein function prediction**

| Organism | Setting | Fmax | | | Accuracy | | |
| --- | --- | --- | --- | --- | --- | --- | --- |
|  |  | BP | MF | CC | BP | MF | CC |
| Human | Gamma_neg:0 (cross-entropy) | 0.447 | 0.204 | 0.392 | 0.363 | 0.032 | 0.210 |
|  | Gamma_neg:1 | 0.459 | 0.187 | 0.436 | **0.368** | 0.029 | 0.294 |
|  | Gamma_neg:2 (DualNetGO) | **0.459** | **0.226** | **0.464** | 0.330 | 0.033 | **0.303** |
|  | Gamma_neg:3 | 0.383 | 0.179 | 0.307 | 0.099 | **0.042** | 0.193 |
| Mouse | Gamma_neg:0 (cross-entropy) | 0.271 | 0.440 | 0.501 | **0.103** | 0.333 | 0.231 |
|  | Gamma_neg:1 | **0.302** | 0.434 | **0.513** | 0.052 | 0.317 | **0.347** |
|  | Gamma_neg:2 (DualNetGO) | 0.296 | **0.524** | 0.502 | 0.090 | **0.374** | 0.224 |
|  | Gamma_neg:3 | 0.288 | 0.397 | 0.485 | 0.077 | 0.294 | 0.197 |

**Table S3.** Hyperparameter search results for different values of gamma_neg of ASL in DualNetGO. The original value is set to 2 in our study, marked by red color. Best results are marked in bold and the second best are underlined.

**5. Effects of different NLP layers and dropout values of the Classifier in DualNetGO**

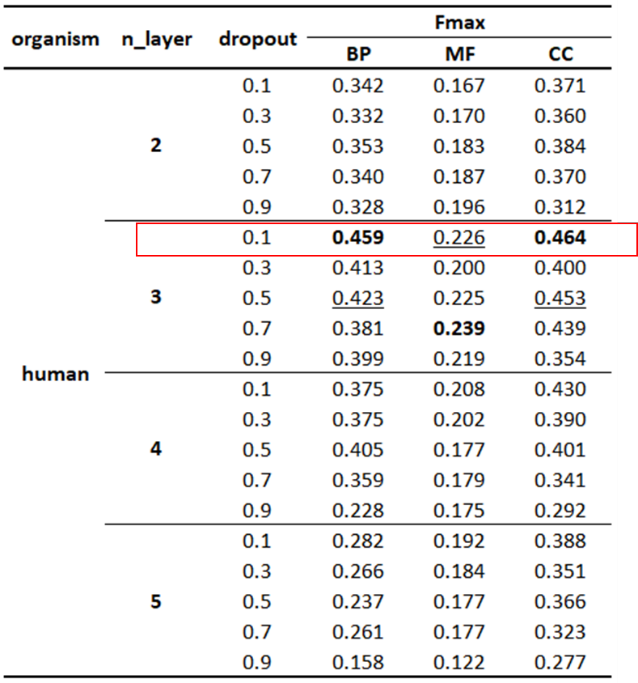

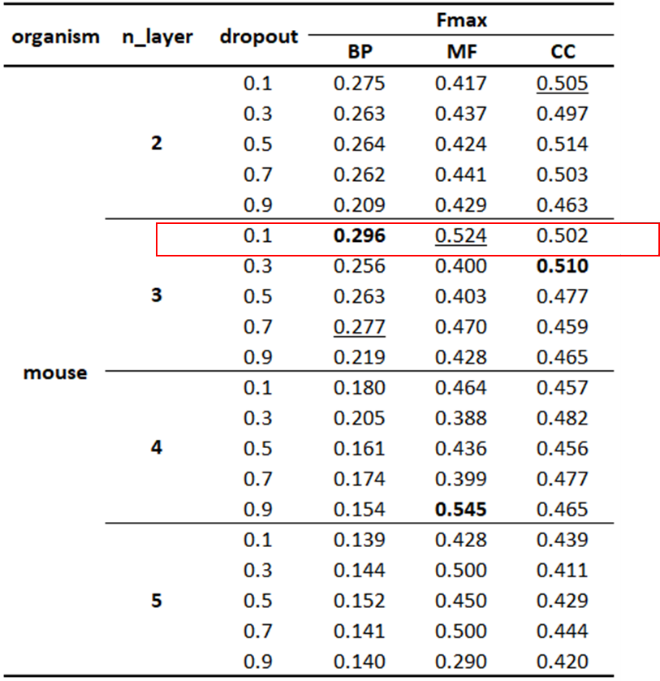

**Table S4.** Hyperparameter search results for different layer numbers and dropouts of the Classifier in DualNetGO. The original value of n_layer is set to 3 and dropout rate set to 0.1 in our study, marked by red rectangles. Best results are marked in bold and the second best are underlined.

**6. Details about DualNetGO model and training parameters**

The training process is divided into three stages, and two networks are trained alternately in the first two stages. The number of training epochs for each stage is denoted as E1, E2, and E3, respectively, and the maximum number of feature matrices to be included for prediction is denoted as N_f_. These numbers are set as hyperparameters before training.

In the first stage, a random combination of feature matrices is sampled from the matrix space $M$ with no more than N_f_ matrices at the beginning of each epoch. These matrices go through the classifier and an ASL prediction loss on validation set L_c_ is calculated. L_c_ is further backpropagated to update weights of corresponding MLP modules in the classifier for each selected matrix. The combination of selected matrices is encoded as a one-hot encoding mask m_1_, and together with L_c_ is used as input for the selector. An MSE loss L_s_ is calculated between the output of the selector and L_c_, and backpropagated to update weights in the selector. In total E1 combinations are evaluated, and weights in the MLP module for each matrix in the classifier are updated. The number of rounds for the weights in an MLP module to be updated depends on how many times the corresponding matrix is sampled.

In the second stage, in each epoch we first create a mask with equal weights of 0.5 representing an equal chance for each matrix to be selected, and then we use this mask as input for the selector and calculate the gradient of each element in the mask with respect to the model. Given a trained selector model, gradients of the input are expected to reflect the importance of each element. We select the corresponding matrices with top N_f_ absolute gradient values, indicated by the indices in the mask, to form a new matrix space M_f_. Because the optimal combination could be a subset of M_f_, 10 combinations are sampled from M_f_ and evaluated by the classifier on the validation set. The lowest loss and the corresponding mask m2 are recorded, and m2 is used for a similar process in stage 1 to train the classifier and then the selector. Different epochs in stage 2 may generate different M_f_s as weights are updated for the two networks. In total, several E2 losses and masks are recorded.

In the third stage, the mask with the minimal validation loss across the record is identified as m*, and the training process for the classifier continues only using the corresponding matrices with m*. Weights in the MLP modules with respect to m* and the prediction head in the classifier are updated for E3 epochs. Performance on test set data is reported as the final results when the classifier achieves the best Fmax score on the validation set.

Theoretically, there are $\sum_{i=1}^{N_{f}} \binom{\left| M \right|}{i}$ combinations. Assuming each combination needs 100 epochs to train and all combinations are enumerated to determine the best one, the total training time will be intractable (given |M|=8 and N_f_=4, the total number of epochs is 16200). However, in DualNetGO only one combination is sampled in each epoch of stage 1, and 10 combinations in each epoch for stage 2. The total number of training epochs is E1+10*E2+E3. According to our possible hyperparameter settings (**Table S5**) the maximum number of weight updates for the classifier is 1410 epochs. We acknowledge that the final subset calculated by DualNetGO may not be the optimal one due to the sampling nature in stage 1 and 2. As training DualNetGO is a stochastic process, a careful choice of the hyperparameters of the epochs in stages 1 and 2 may be necessary to approach the optimal solution, as suggested by another dual-network model that deals with heterophilic graph data. Fortunately, the search of hyperparameters costs little time, and DualNetGO is more efficient in determining a suitable combination of features than enumerating all possibilities.

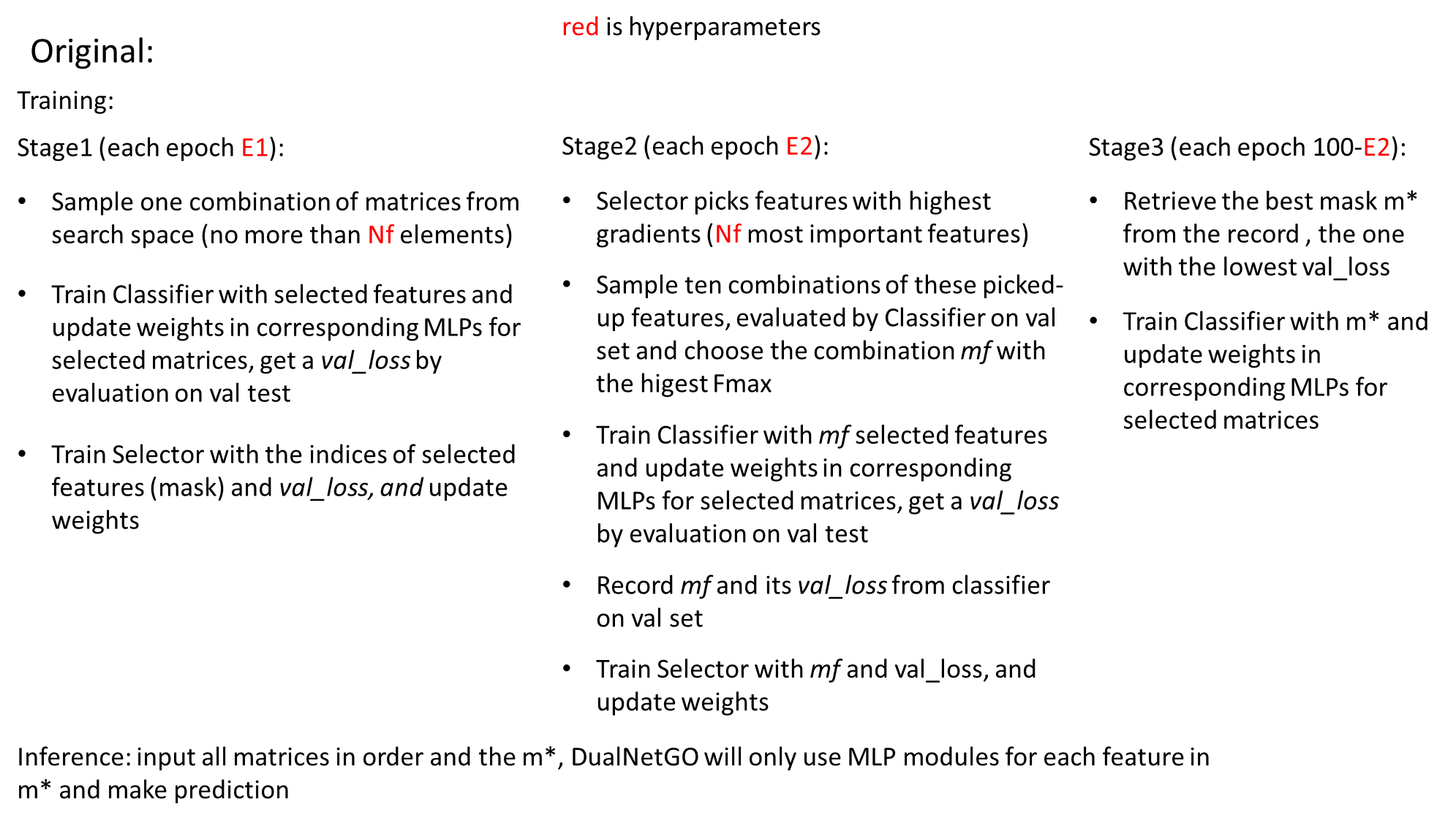

For DualNetGO, then number of hidden dimensions of the classifier is set to 512 except for the prediction head whose dimension is set to the number of GO terms, and 256 for the selector. Dropout rates are set to 0.5 for all components except for the prediction head, which is set to 0.1. Learning rates are set to 0.01 for both the classifier and the selector. AdamW is chosen as the optimizer for the classifier and the selector with weight decays set to 0 and 1e-5, respectively. Hyperparameters are determined by manual grid search on the validation data (**Table S5**). All experiments are conducted with a single RTX 3090 GPU with 24G memory. A single run for training DualNetGO takes 2-5 minutes and 2-6 GB GPU memory depending on the number of iterations and size of training data.

| **Hyperparameters** | **Search Space** |
| --- | --- |
| Iteration epoch for stage1 | [100, 200, 300, 400, 500] |
| Iteration epoch for stage2 | [10, 20, 30, 40, 50, 60, 70, 80, 90] |
| Maximum number of features for selection | [2, 3, 4, 5] |

**Table S5.** Model hyperparameter search space. Other possible hyperparameters include learning rate for classifier and selector, dropout for linear layer output, dimension for hidden states, and weight decay for Adam optimizer.

**7. Concept of embedding-centric versus evidence-centric DualNetGO model**

We denote the previous search space for searching across different PPI networks given a specific graph embedding method (TransformerAE) as the **embedding-centric setting**. We call the method of utilizing DualNetGO model the **evidence-centric setting**, which searches across different graph embedding algorithms given a PPI network from a specific type of evidence. Differences between the embedding-centric and evidence-centric setting of DualNetGO lie in the feature matrices included in the search space as shown in **Figure S1**. In the embedding-centric setting, one graph embedding method is chosen and used to preprocess different PPI networks. The encoded graph embeddings are included in the search space. In the evidence-centric setting, one type of PPI network is chosen and different graph embedding methods are used for preprocessing. As an extension to DualNetGO, we also implement the model in the evidence-centric setting using only the combined PPI network and evaluate the performance on both human and mouse datasets. Collectively, DualNetGO with embedding-centric and evidence-centric settings produces the top-2 results in almost all metrics in all datasets except in mouse MF where Graph2GO generally performs better (**Table S6**). We also evaluate the performances of using other graph embedding methods for the embedding-centric model and other PPI networks for the evidence-centric model. Both yield good results. The best performance and the corresponding parameters can be found in **Table S10, S11**.

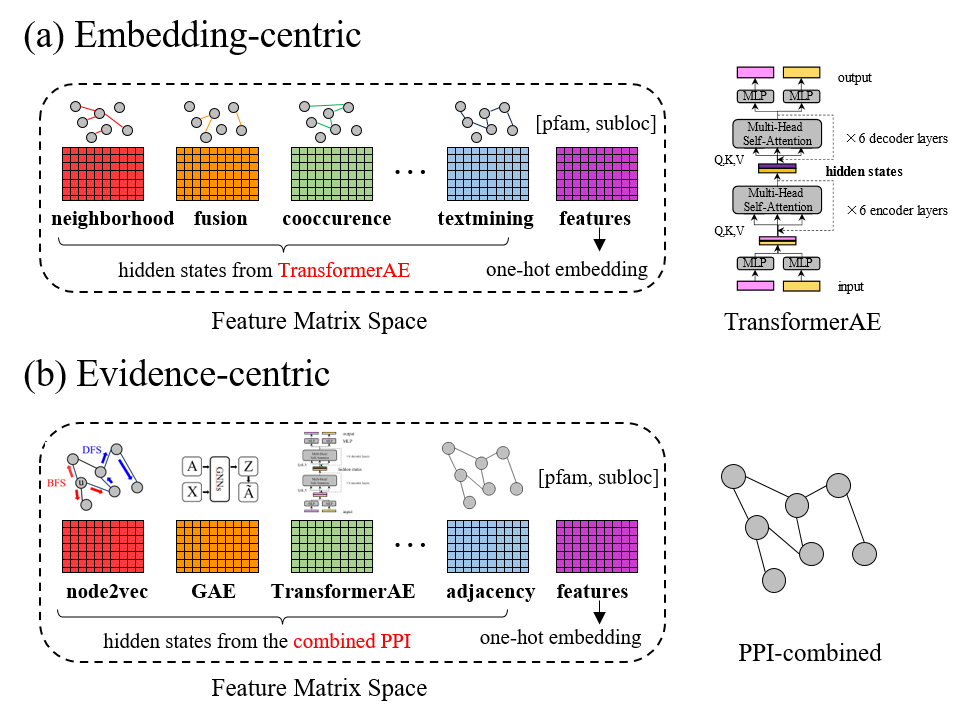

**Figure S1.** Comparison of Embedding-centric setting and evidence-centric setting.

**8. Detailed results of DualNetGO with embedding-centric and evidence-centric setting and corresponding parameters**

Detailed metrics corresponding to **Figure 3** in the paper are shown in **Table S6**. TransformedAE is fixed for the embedding-centric model and combined PPI is fixed for the eveidence-centric model in this section.

| Organism | Model | BP | | | | MF | | | | CC | | | |
| --- | --- | --- | --- | --- | --- | --- | --- | --- | --- | --- | --- | --- | --- |
|  |  | M-AUPR | m-AUPR | Accuracy | Fmax | M-AUPR | m-AUPR | Accuracy | Fmax | M-AUPR | m-AUPR | Accuracy | Fmax |
| Human  (9606) | Naïve | 0.048 | 0.024 | 0.000 | 0.051 | 0.029 | 0.032 | 0.000 | 0.099 | 0.059 | 0.049 | 0.000 | 0.087 |
|  | BLAST | 0.046 | 0.029 | 0.000 | 0.067 | 0.036 | 0.028 | 0.003 | 0.041 | 0.057 | 0.034 | 0.000 | 0.059 |
|  | Mashup | 0.165 | 0.260 | **0.346** | 0.395 | 0.111 | 0.082 | 0.049 | 0.130 | 0.151 | 0.162 | 0.092 | 0.280 |
|  | deepNF | 0.127 | 0.218 | 0.154 | 0.331 | 0.088 | 0.067 | 0.045 | 0.116 | 0.135 | 0.163 | 0.067 | 0.221 |
|  | Grap2GO | 0.144 | 0.167 | 0.187 | 0.289 | 0.100 | 0.084 | 0.014 | 0.172 | 0.138 | 0.149 | 0.050 | 0.188 |
|  | CFAGO | 0.184 | 0.306 | 0.253 | 0.414 | **0.126** | 0.107 | **0.060** | 0.164 | 0.193 | 0.281 | 0.160 | 0.322 |
|  | DualNetGO  (embedding) | **0.204** | 0.364 | 0.330 | 0.459 | 0.117 | **0.133** | 0.033 | 0.226 | **0.256** | 0.422 | **0.303** | **0.464** |
|  | DualNetGO  (evidence) | 0.199 | **0.385** | 0.225 | **0.483** | 0.118 | 0.128 | 0.004 | **0.245** | 0.198 | 0.392 | 0.116 | 0.443 |
| Mouse  (10090) | Naïve | 0.051 | 0.035 | 0.000 | 0.085 | 0.064 | 0.058 | 0.000 | 0.171 | 0.045 | 0.124 | 0.000 | 0.192 |
|  | BLAST | 0.051 | 0.035 | 0.000 | 0.059 | 0.067 | 0.059 | 0.008 | 0.066 | 0.047 | 0.055 | 0.007 | 0.134 |
|  | Mashup | 0.161 | 0.148 | 0.052 | 0.247 | 0.230 | 0.281 | 0.222 | 0.365 | 0.135 | 0.230 | 0.143 | 0.301 |
|  | deepNF | 0.097 | 0.095 | 0.013 | 0.155 | 0.205 | 0.295 | 0.183 | 0.335 | 0.090 | 0.184 | 0.088 | 0.280 |
|  | Grap2GO | 0.108 | 0.101 | 0.000 | 0.195 | **0.254** | 0.490 | **0.492** | **0.595** | 0.148 | 0.280 | 0.231 | 0.348 |
|  | CFAGO | **0.187** | 0.185 | 0.071 | 0.269 | 0.252 | 0.374 | 0.167 | 0.418 | 0.201 | 0.352 | **0.224** | 0.425 |
|  | DualNetGO  (embedding) | 0.155 | **0.186** | **0.090** | 0.296 | 0.207 | 0.460 | 0.373 | 0.524 | 0.229 | **0.459** | **0.224** | **0.502** |
|  | DualNetGO  (evidence) | 0.174 | 0.172 | 0.084 | **0.296** | 0.229 | **0.504** | 0.222 | 0.558 | **0.242** | 0.434 | 0.150 | 0.478 |

**Table S6.** Performance of DualNetGO with detailed metrics. Best performance numbers are shown in bold and the second best are underlined.

The hyperparameters for attaining our results can be found in **Table S7**.

| Organism | Setting | Aspect | E1 | E2 | E3 | $N_{f}$ |
| --- | --- | --- | --- | --- | --- | --- |
| Human  (9606) | TransformerAE  (embedding) | BP | 300 | 10 | 90 | 4 |
|  |  | MF | 100 | 40 | 60 | 4 |
|  |  | CC | 500 | 80 | 20 | 3 |
|  | combined  (evidence) | BP | 500 | 80 | 20 | 4 |
|  |  | MF | 200 | 10 | 90 | 3 |
|  |  | CC | 100 | 50 | 50 | 4 |
| Mouse  (10090) | TransformerAE  (embedding) | BP | 100 | 90 | 10 | 4 |
|  |  | MF | 100 | 90 | 10 | 4 |
|  |  | CC | 400 | 10 | 90 | 4 |
|  | combined  (evidence) | BP | 300 | 60 | 40 | 3 |
|  |  | MF | 400 | 80 | 20 | 2 |
|  |  | CC | 200 | 30 | 70 | 2 |

**Table S7.** Hyperparameters for DualNetGO with fixed settings.

**9. Comprehensive comparison among different settings of PPI on mouse**

We also conduct experiments on the mouse dataset across different graph embedding methods and different evidence. Results show that DualNetGO outperforms models with other graph embedding methods.

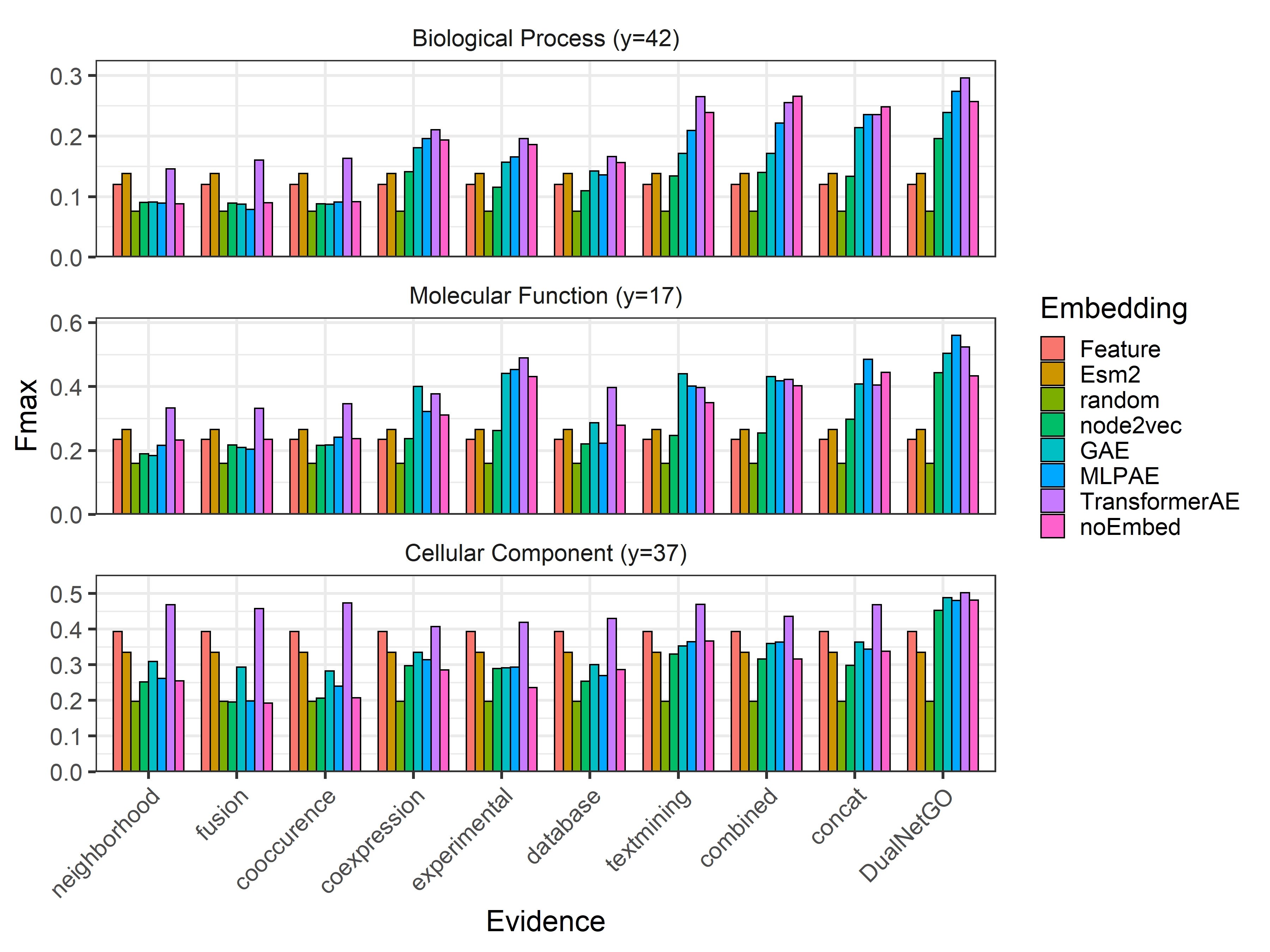

**Figure S2.** Comparison of different graph embedding methods on different PPI networks of mouse.

**10. Designs of ablation tests on the Selector, the Stage 1 and Stage 2 of DualNetGO**

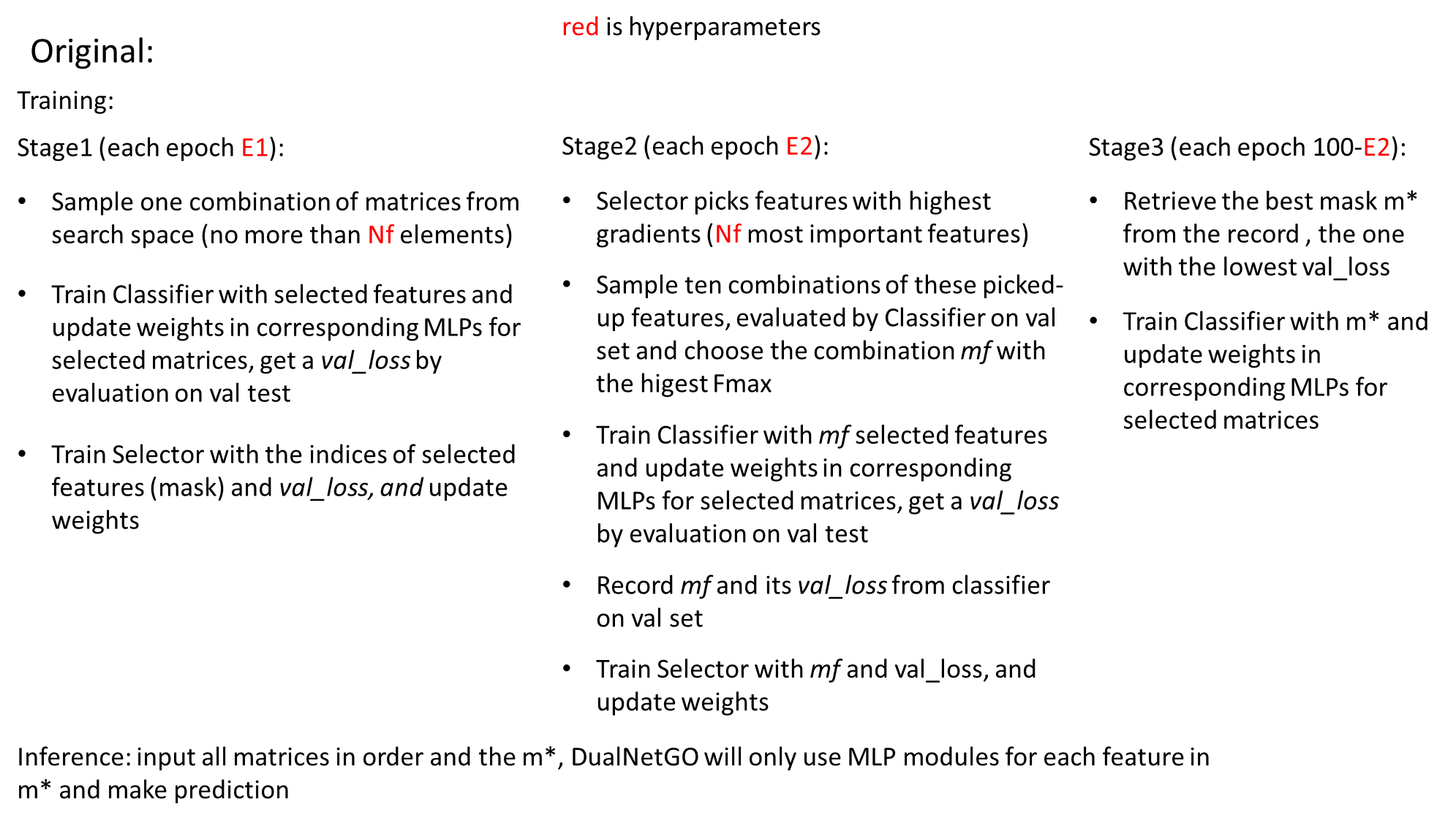

In the original DualNetGO model, Stage 1 serves as an exploration process to gather information of performances of different subsets by training a Selector using the feature mask and the corresponding validation loss. Stage 2 serves as an exploitation process to give a more accurate evaluation on the importance of each feature matrix and further narrow down the possible choices of feature combinations.

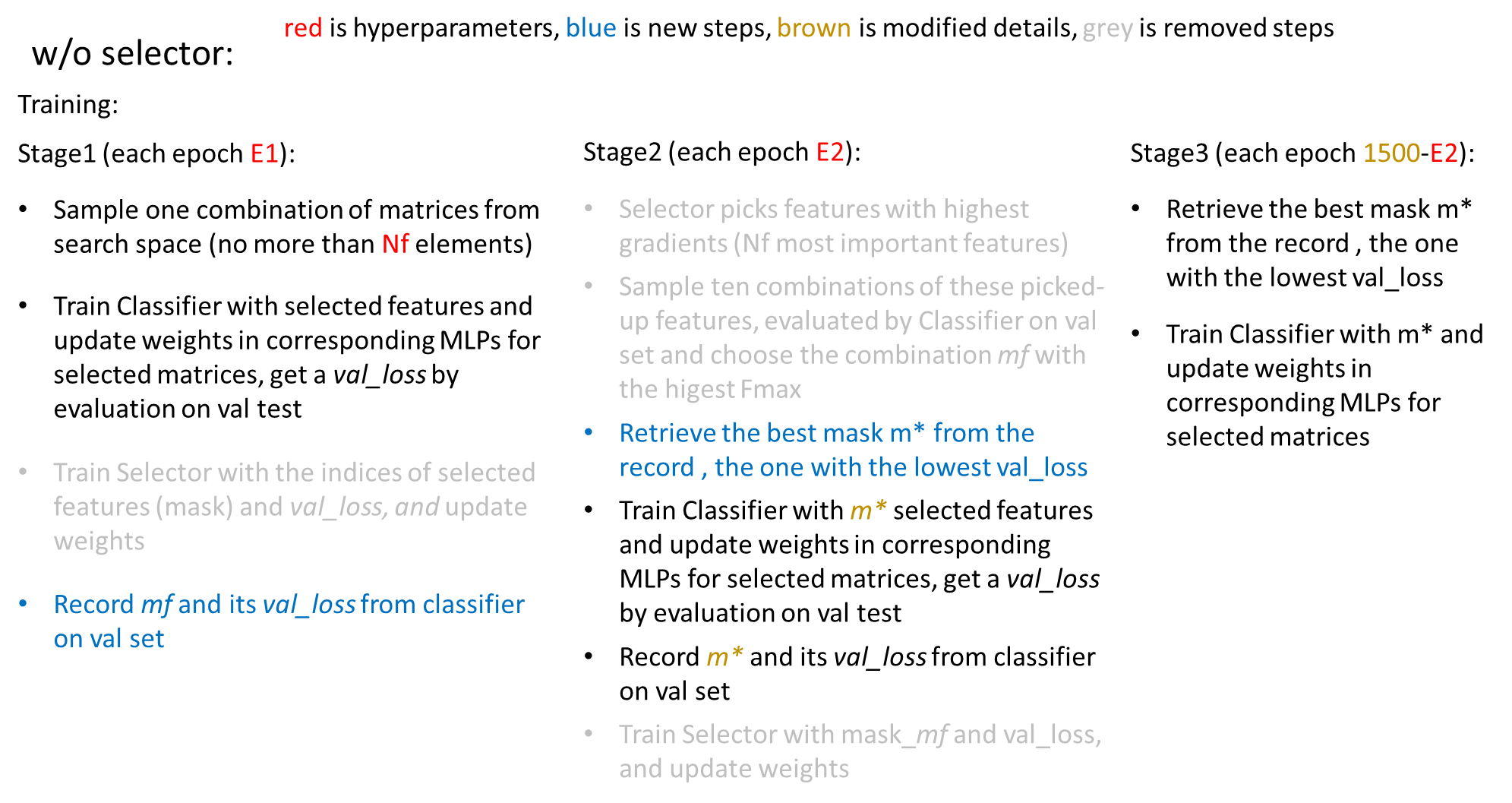

The Selector evaluates the importance of each feature matrix by extracting the corresponding gradients with respect to the input. Without Selector, the selection of the best features completely depends on the previously recorded performance of the Classifier on the validation set. In each epoch (E1) only a subset of feature matrices is selected and only the weights of the corresponding MLP modules are updated, resulting in a stochastic training process. However, Stage 1 cannot guarantee that every combination is encountered, and even if a specific combination is trained, the training is only of a couple of epochs. In contrast, Selector can give a relatively fair evaluation of all feature matrices without testing all combinations after the exploration process of Stage 1.

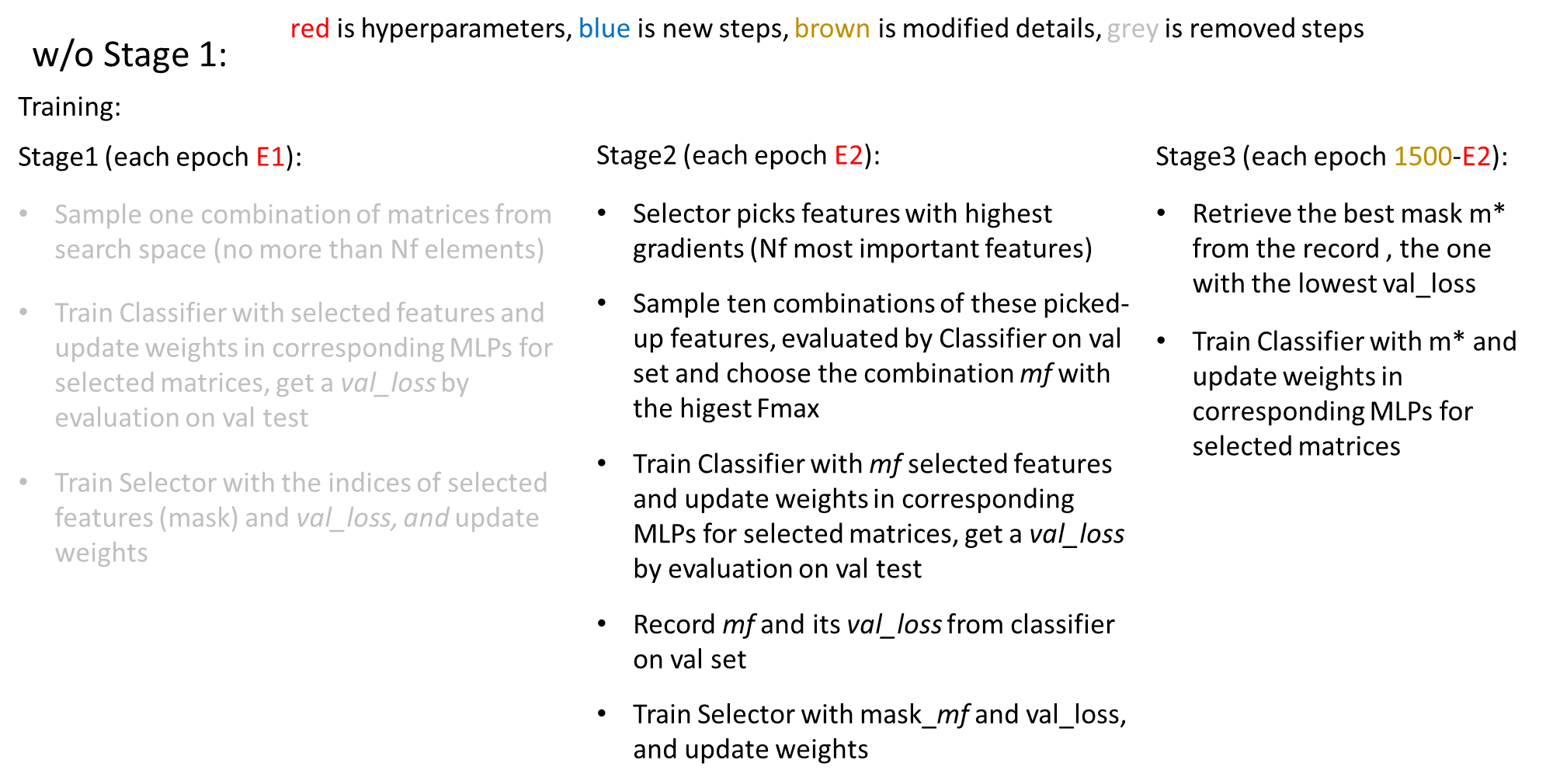

As an exploration process, Stage 1 gathers information for the Selector to accurately evaluate the importance of each feature matrix in Stage 2. Since the evaluation is based on the gradients of the Selector model, without Stage 1 the Selector will be randomly initiated, and thus the gradients have nothing to do with feature importance without the prior knowledge gathered in Stage 1 through Classifier’s performance on the validation set. With a large number of epochs in Stage 2 (E2), the Selector may gradually produce accurate evaluation as more and more combinations are sampled, but is not as efficient as that in Stage 1, because the combinations sampling in Stage 2 is not fully random but depends on the Selector.

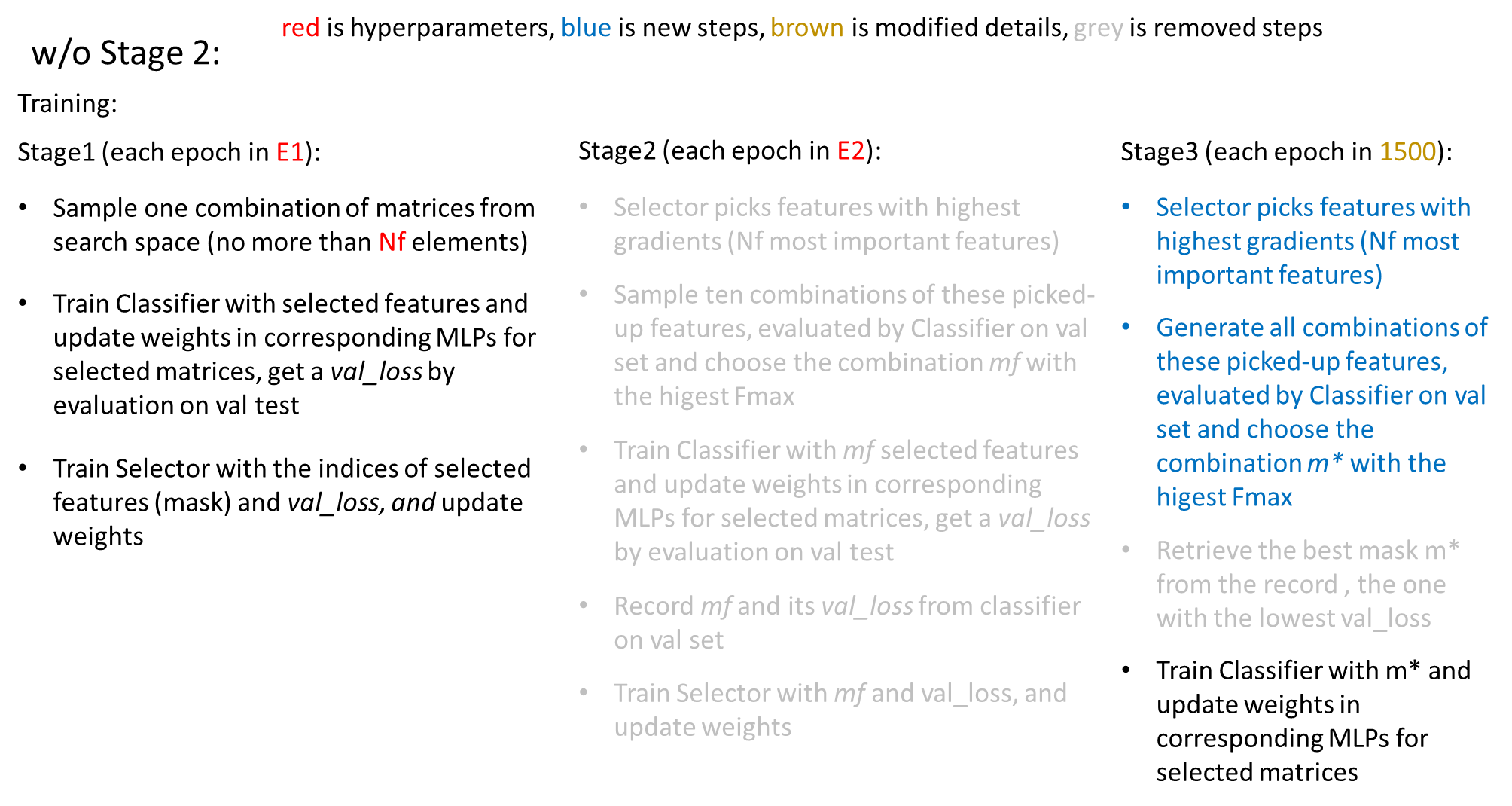

In contrast, Stage 2 serves as an exploitation process for the Selector to further narrow down the possible choices of combinations, select one out of ten sampled combinations (no weight updates) with the highest Fmax, and then use the selected mask to further train the Classifier and the Selector (weigh updates). In our experiments, Stage 2 takes 11 inference rounds to further update the model weights, so it is slower than Stage 1 but the mask generated is closer to the optimal one.

**11. Effects of training DualNetGO without the Selector and using all feature matrices as input**

| Organism | Setting | Fmax | | | Accuracy | | |
| --- | --- | --- | --- | --- | --- | --- | --- |
|  |  | BP | MF | CC | BP | MF | CC |
| Human | Original | **0.459** | **0.226** | **0.464** | **0.330** | 0.033 | **0.303** |
|  | All_matrices  separated | 0.236 | 0.143 | 0.263 | 0.132 | **0.050** | 0.101 |
|  | All_matrices  concat | 0.286 | 0.154 | 0.259 | 0.187 | 0.005 | 0.134 |
|  | CFAGO-concat | 0.332 | 0.157 | 0.379 | 0.187 | 0.031 | 0.218 |
| Mouse | Original | **0.296** | **0.524** | **0.502** | **0.090** | **0.374** | 0.224 |
|  | All_matrices  separated | 0.069 | 0.474 | 0.455 | 0.000 | 0.000 | 0.122 |
|  | All_matrices  concat | 0.218 | 0.417 | 0.444 | 0.045 | 0.048 | 0.265 |
|  | CFAGO-concat | 0.265 | 0.479 | 0.467 | 0.084 | 0.349 | 0.170 |

**Table S8.** Ablation test of DualNetGO when training all matrices together and no sampling, and no training for the Selector. The original DualNetGO is marked by red color. Best results are marked in bold and the second best are underlined. **All_matrices** **separated**: 8 MLP modules, each for one matrix; **All_matrices** **concat**: 1 MLP module for a concatenated matrix from all matrices; **CFAGO-concat**: use CFAGO prediction head structure with the DualNetGO training procedure.

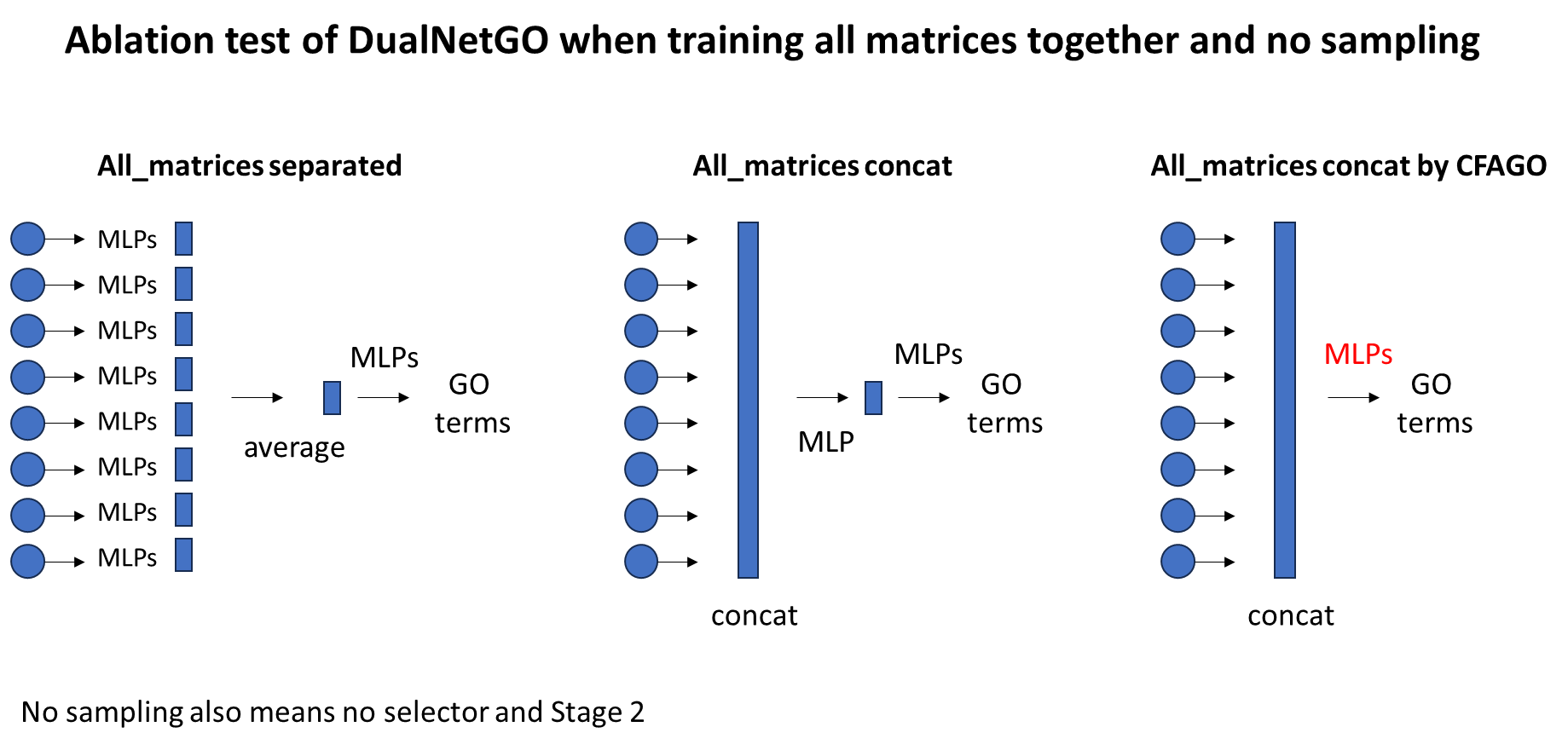

**Figure S3.** An illustration about the different training strategies for training Classifier with all matrices as input without the Selector. The red ‘MLPs’ on the right represents a different MLP prediction head structure used in the CFAGO paper for utilizing embeddings from TransformerAE.

**12. Impacts of different E1, E2 and num_feat values on the performance of DualNetGO**

We provide Fmax scores for different E1, E2, and N_f_ combinations of DualNetGO on human as an example. Results show trends for the changes of Fmax when hyperparameters change gradually. We can see that for BP and MF the Fmax values reduce and CC increase with E1 varies from 100 to 500. The trend depends on the distribution and complexity of the original data. Difficult samples may require a longer process of exploration in Stage 1 but simple data may be overfitted by a larger number of E1. When fixing E1 to investigate the effect of E2, we find that when the model is not overfitted in Stage 1, Fmax scores increase as E2 increases, such as P-E1-100 and those in C; while trends of decrease appear in P-E1-400/500, F-E1-500. One can find the trend through some simple evaluations with a couple of hyperparameters and set values accordingly to train a DualNetGO model with their data.

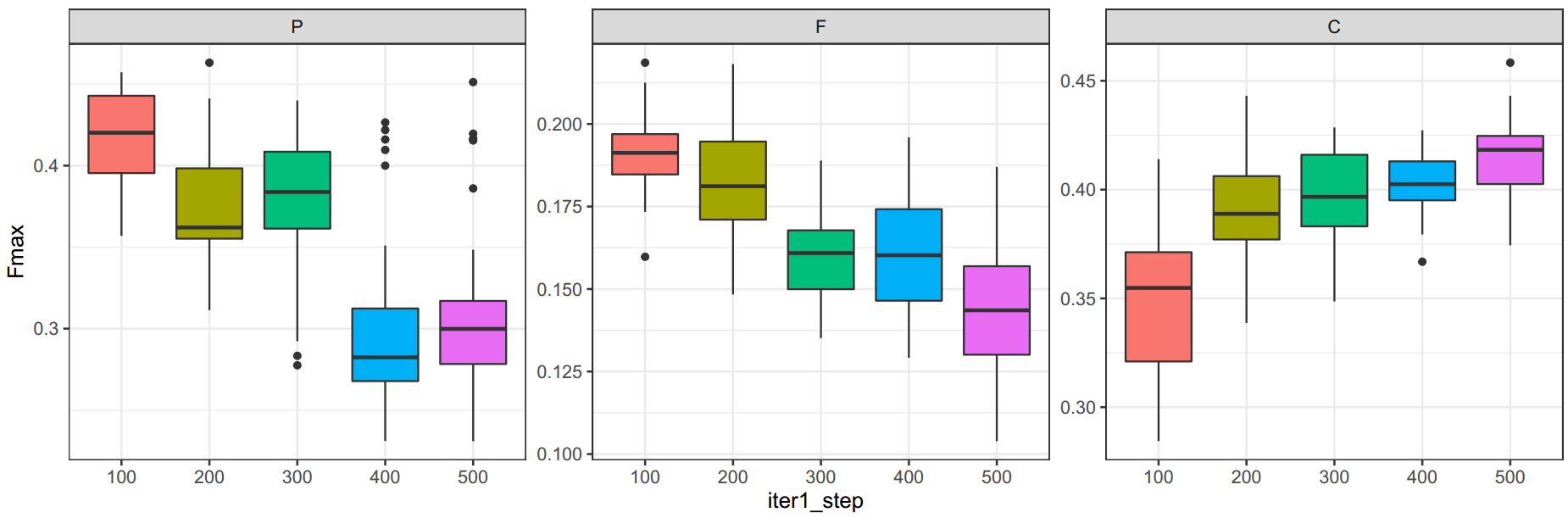

**Figure S4.** Fmax scores on different E1 across different E2 and Nf.

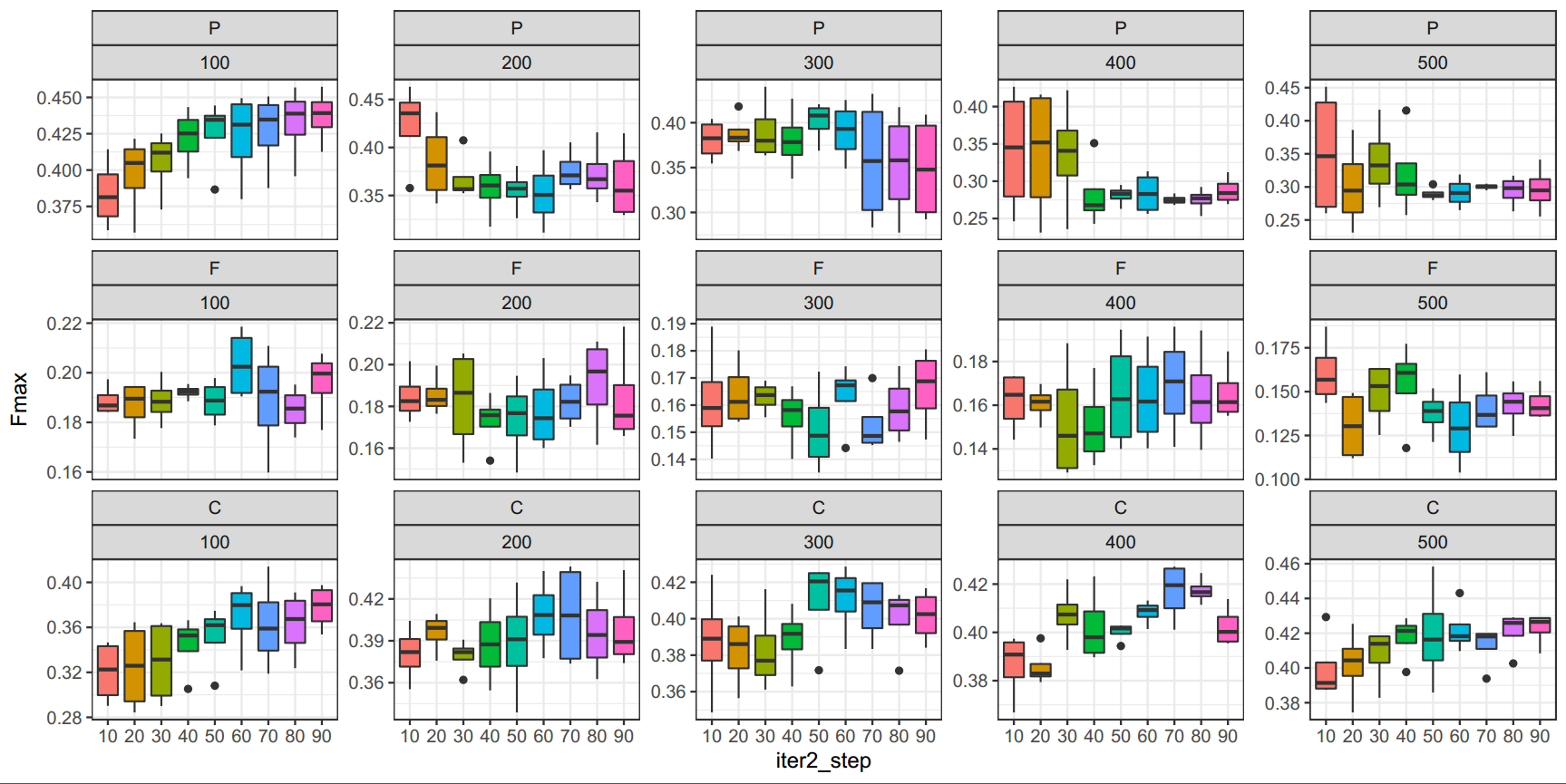

**Figure S5.** Fmax scores on different E1 and E2 across different Nf. Subfigures are grouped according to GO aspects and iter1_step (E1) numbers. The x-axis is the iter2_step (E2) number.

**13. Graph processing time, model training time, and memory usage**

In addition to preprocessing and training time, we also measure the model size and the maximum memory usage for each model included in the study (**Table S9**). DualNetGO with the evidence-centric setting is not specified in this table as it is similar to that with the embedding-centric setting.

| Model | Strategy | Preprocessing (P) | Training (T) | Parameters (P+T) | Max memory |
| --- | --- | --- | --- | --- | --- |
| Mashup | rwr+svd | 1955s | 168s  (svm_512d) | / | 12G (CPU, whole matrix) |
| deepNF | MLPAE | 462s (20 epochs) | 190s (svm_1200d) | 1015M+0M | 19G (batch256) |
| Graph2GO | GAE | 3.78h (blast) +  1437s (GAE 100 epochs) | 21s (mlp 100 epochs) | 0.88M+2.3M | 5G (GAE batch128) |
| CFAGO | TransformerAE | ~50h (5000 epochs) | 710s (finetune-mlp 100 epochs) | 82M+0.27M | 3G (batch32) |
| DualNetGO | node2vec (rwr) | 62s (20 epochs) | 113s (mlp > 200 epochs) | 10M+2.8M | 5G (node2vec batch128) |
|  | MLPAE | 866s (100 epochs) |  | 41M+2.8M | 2.5G (MLPAE batch32) |
|  | GAE | 323s (20 epochs) |  | 2.1M+2.8M | 4G (GAE batch128) |
|  | NoEmbed | <1s |  | 0M+70M | 6G |
|  | TransformerAE | ~50h (5000 epochs) |  | 82M+2.8M | 3G (batch32) |

**Table S9.** Details of elapsed time and memory usage for different models.

**14. Best Fmax scores of DualNetGO with embedding-centric and evidence-centric setting and corresponding parameters**

The performance of DualNetGO improves when evaluated across all evidence and all graph embedding methods, but not in a mixed setting where different PPI networks and graph embedding methods co-exist at the same time in the feature matrix space for search. Here we show the best performance produced by DualNetGO in comparison with a previous SOTA model CFAGO (**Table S10**).

| Organism | | Model | BP | | | | | MF | | | | | CC | | | | |
| --- | --- | --- | --- | --- | --- | --- | --- | --- | --- | --- | --- | --- | --- | --- | --- | --- | --- |
|  |  |  | Setting | M-AUPR | m-AUPR | Acc. | Fmax | Setting | M-AUPR | m-AUPR | Acc. | Fmax | Setting | M-AUPR | m-AUPR | Acc. | Fmax |
| Human  (9606) | CFAGO | | / | 0.184 | 0.306 | 0.253 | 0.414 | / | 0.126 | 0.107 | **0.060** | 0.164 | / | 0.193 | 0.281 | 0.160 | 0.322 |
|  | DualNetGO  (embedding) | | No Embedding | 0.180 | 0.346 | **0.407** | 0.468 | MLP-AE | **0.129** | **0.137** | 0.029 | 0.235 | Trans-formerAE | **0.256** | 0.422 | 0.303 | 0.464 |
|  | DualNetGO  (evidence) | | com-bined | **0.199** | **0.385** | 0.225 | **0.483** | com-bined | 0.118 | 0.128 | 0.004 | **0.245** | neigh-borhood | 0.254 | **0.493** | **0.361** | **0.502** |
| Mouse  (10090) | CFAGO | | **/** | **0.187** | 0.185 | 0.071 | 0.269 | / | 0.252 | 0.374 | 0.167 | 0.418 | / | 0.201 | 0.352 | 0.224 | 0.425 |
|  | DualNetGO  (embedding) | | Trans-formerAE | 0.155 | **0.186** | **0.090** | 0.296 | MLP-AE | 0.229 | 0.459 | 0.224 | 0.502 | Trans-formerAE | **0.229** | **0.459** | 0.224 | 0.502 |
|  | DualNetGO  (evidence) | | com-bined | 0.174 | 0.172 | 0.084 | **0.296** | text-mining | **0.244** | **0.493** | **0.452** | **0.581** | neigh-borhood | 0.227 | 0.436 | **0.286** | **0.504** |

**Table S10.** Best Fmax performance of DualNetGO with corresponding settings. Highest scores are in bold.

The corresponding hyperparameters to attain these results are shown in **Table S11**.

| Organism | Model | Setting | Aspect | E1 | E2 | E3 | $N_{f}$ |
| --- | --- | --- | --- | --- | --- | --- | --- |
| Human  (9606) | Embedding-centric | NoEmbed | BP | 400 | 70 | 30 | 3 |
|  |  | MLPAE | MF | 500 | 40 | 60 | 2 |
|  |  | TransformerAE | CC | 500 | 80 | 20 | 3 |
|  | Evidence-centric | combined | BP | 500 | 80 | 20 | 4 |
|  |  | combined | MF | 200 | 10 | 90 | 3 |
|  |  | neighborhood | CC | 500 | 40 | 60 | 5 |
| Mouse  (10096) | Embedding-centric | TransformerAE | BP | 100 | 90 | 10 | 4 |
|  |  | MLPAE | MF | 100 | 90 | 10 | 5 |
|  |  | TransformerAE | CC | 400 | 10 | 90 | 4 |
|  | Evidence-centric | combined | BP | 300 | 60 | 40 | 3 |
|  |  | textmining | MF | 200 | 70 | 30 | 2 |
|  |  | neighborhood | CC | 500 | 50 | 50 | 2 |

**Table S11**. Hyperparameters for DualNetGO with best Fmax scores.

**15. Data Collection for 20 species in CAFA3 dataset**

**A. PPI information from the STRING database**

As stated in the manuscript, the processed CAFA3 dataset can be downloaded from the TEMPROT paper (Oliveira et.al., 2023). We first manually retrieve the taxonomy code of each protein in the CAFA3 test set using the first seven digits of the provided IDs from the CAFA3 competition. For example, given that a CAFA3 ID is “T100900000026”, the first seven digits are 1009000. We print out all len-7 numbers and manually refine them with the current knowledge of NCBI taxonomy codes. Specifically, we remove all 0s at the end of each len-7 number except for 10090 (mouse) which ends with a 0. As a result, we obtain 20 different species from the CAFA3 test set on the BP and MF aspect, and 18 species on the CC aspect. In total, we use the 20 species for downstream training. Although the CAFA3 training and validation sets may contain more species than the 20 species we use, we confine the training set to the 20 species in the test set to reduce the workload of data collection and computational cost of model training, which could result in reduced performance. The 20 species collected are shown in **Table S12**.

We also use PPI information recorded in the STRING database. The experiments conducted on the previous filtered human/mouse UniProt dataset directly used data provided by the CFAGO paper (Wu et al., 2023). However, we found that both the UniProt and STRING databases had significant updates. Some arrangements of the features are different from those provided by CFAGO and thus they cannot be preprocessed using the original script. We have provided a new script to preprocess the new UniProt data for training DualNetGO on CAFA3 data. In addition, the UniProt database updates every eight weeks so the current version has used the new STRING IDs provided in STRING v12.0. The STRING IDs in STRING v11.5 are no longer consistent with those recorded in the latest UniProt. For this reason, we need to use STRING v12.0 to collect PPI information for training and testing DualNetGO under the CAFA3 multi-species setting. Comparing STRING v12.0 (**Table S12**) with v11.5 (**Table S13**), we find that the new STRING database now contains significantly more proteins than those in the older version, making the PPI network information more and more valuable and suitable for general protein properties prediction.

Specifically, given a taxonomy code, we filter the STRING dataset in the “DOWNLOAD” site with the corresponding **species name**, eg. Homo sapiens for 9606 (**Figure S6**). Then we download three files: ***.protein.links.detailed.v12.0.txt.gz**, ***.protein.info.v12.0.txt.gz and** ***.protein.squences.v12.0.fa.gz**. Note that the taxonomy code in STRING can be different from those retrieved in the CAFA test data. In **Table S13**, we put taxonomy codes used in STRING in the parathesis, and those used in CAFA in front of the parathesis. Taxonomy codes used in this study are also provided in **Table S12**.

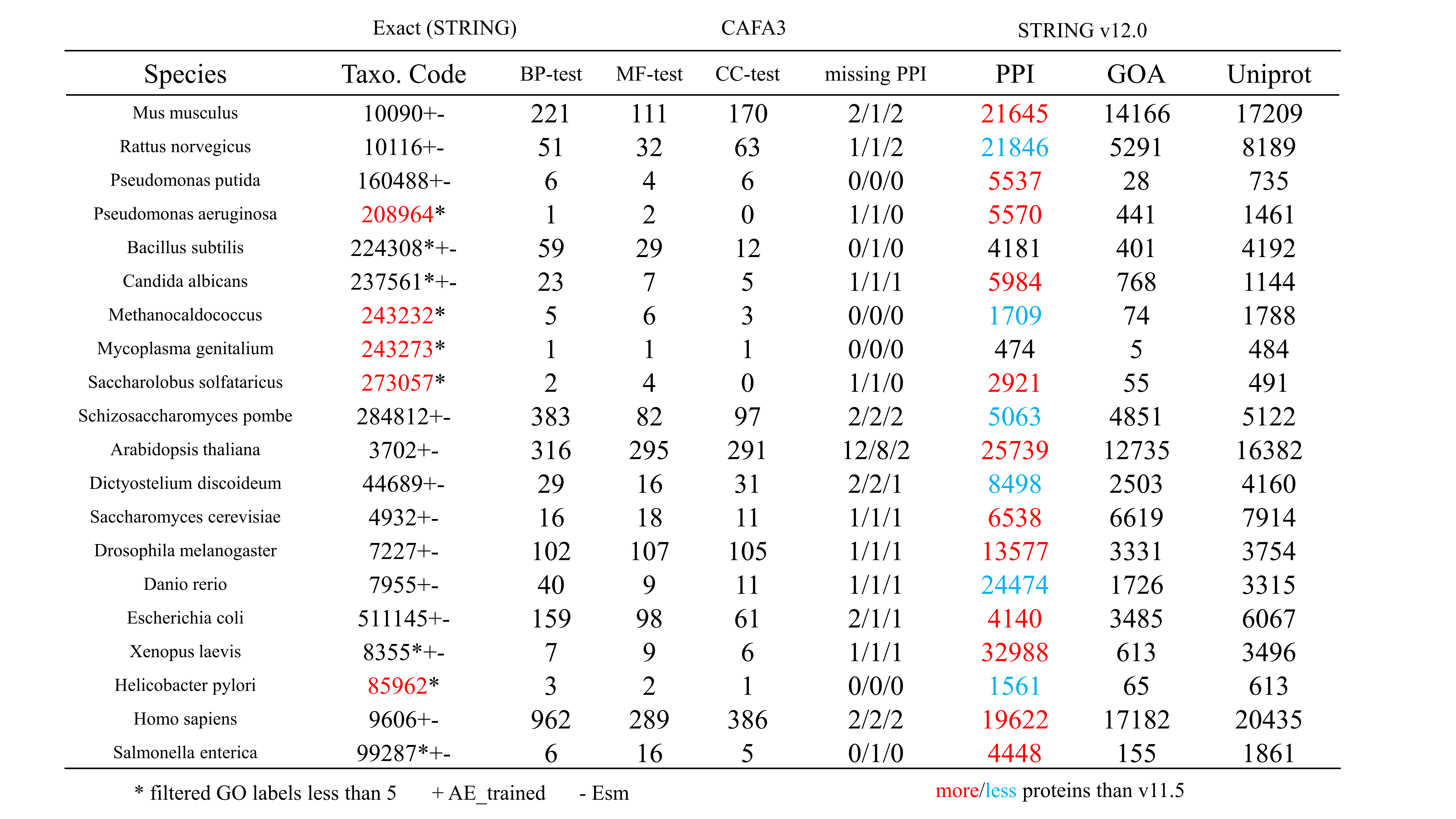

**Table S12.** Statistics of 20 species collected in the CAFA3 test set. Total numbers of proteins in BP, MF and CC test set are 2392, 1137, and 1265, respectively. “missing PPI” denotes the number of CAFA3 test set proteins in the corresponding aspect that lack information in the STRING (v12.0) PPI database. “PPI” denotes the number of proteins present in the PPI network for a certain species. “GOA” denotes the number of proteins that have GO annotations from the QuickGO website. “UniProt” denotes the number of proteins that have reviewed Pfam/subcellular_location information in the UniProt database. Taxonomy with * means that species have few functional annotations and is not suitable for training a separate single-species prediction model. Taxonomy with +/- means the corresponding hidden states of PPIs or Esm2 sequence embeddings are stored and provided on zenodo and can be retrieved. Species in red are not included in the training set because they have few proteins in the test set and few attributes recorded in UniProt.

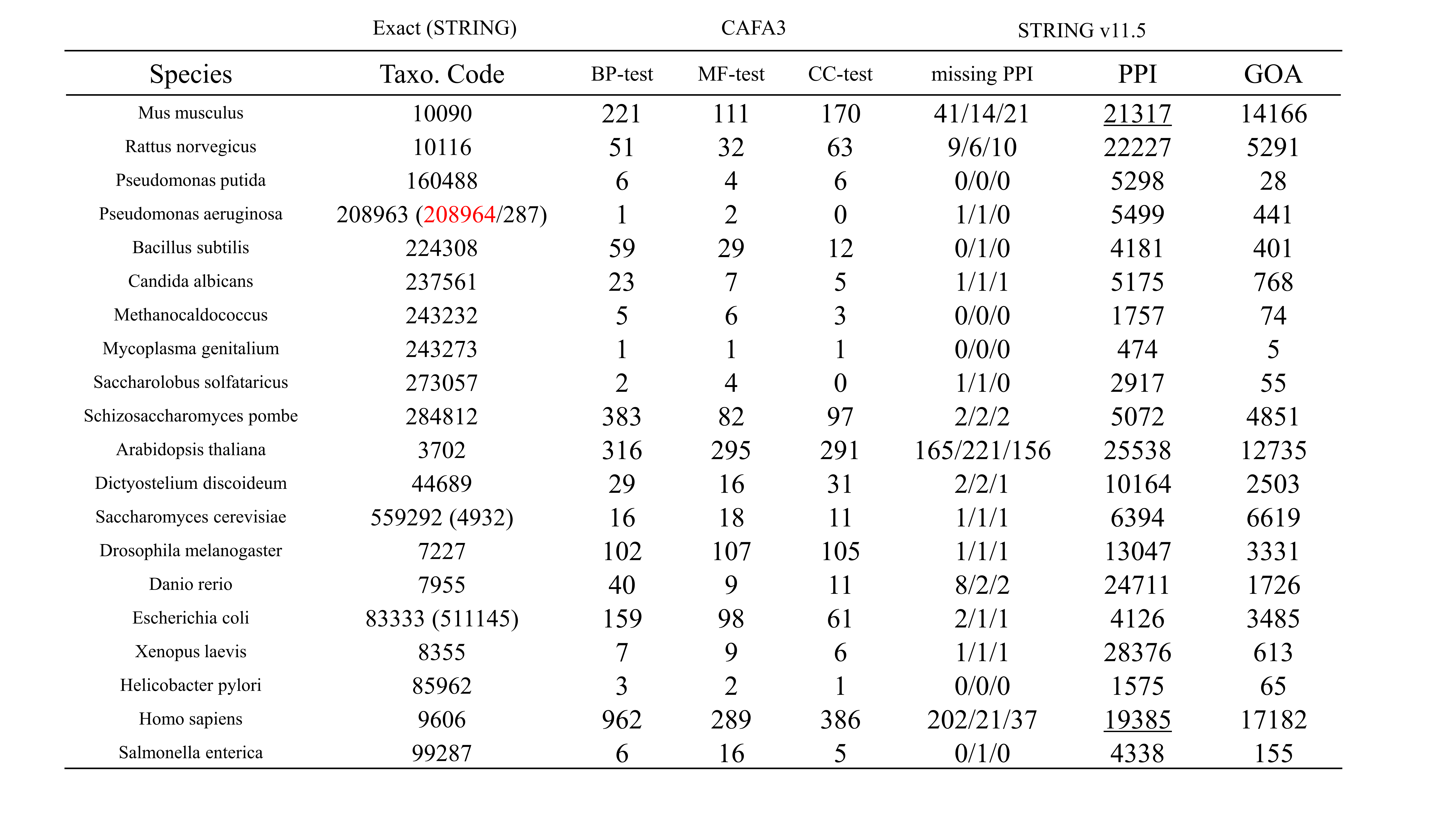

**Table S13.** Statistics of 20 species collected in the CAFA3 test set in an older version of STRING (v11.5). The table shows that it significantly lacks more proteins in the CAFA3 test data set than the newest version. Some of the STRING IDs in this version are deprecated and no longer used by the UniProt database.

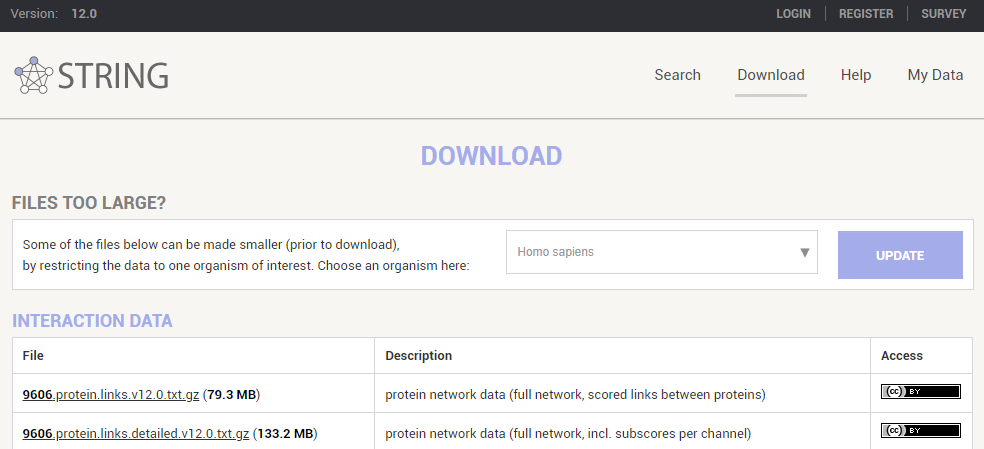

species name

SRTING version

Taxo. code

**Figure S6**. Download site of STRING and settings for gathering PPI network information. Three files are downloaded for each species in total, which are *.protein.links.detailed.v12.0.txt.gz, *.protein.info.v12.0.txt.gz and *.protein.squences.v12.0.fa.gz. Important settings are marked by red rectangles. Species can only be chosen by using the species name.

**B. Annotation for Pfam and subcellular location in the UniProt database**

At the UniProtKB query site, we first filter the database by the corresponding taxonomy code (those included in **Table S12**), and then only retain those that have been reviewed. An example is shown in **Figure S7**.

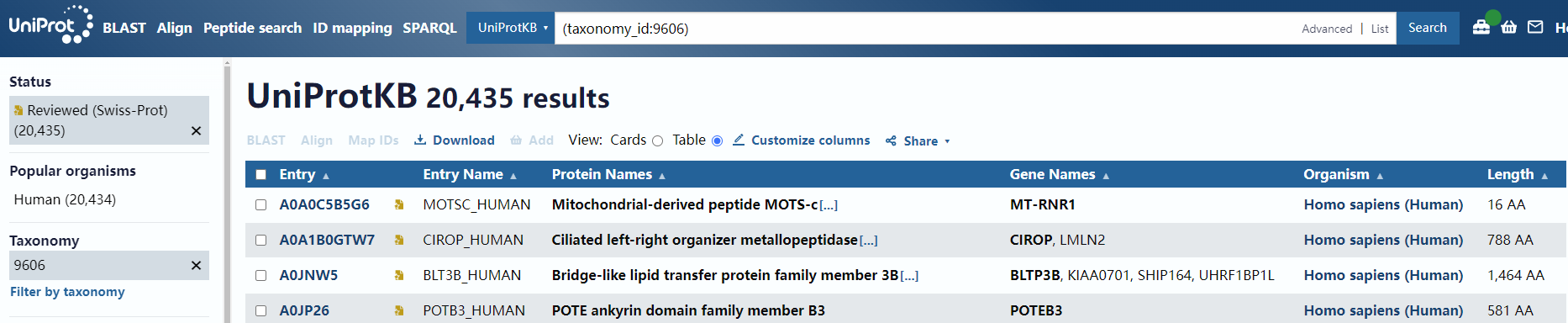

click download after filtering

**Figure S7**. Query site of UniProt database and setting for gathering protein information. Filtering options are marked by red rectangles. Taxonomy can be set by taxonomy code.

After filtering, we click the download icon and choose the TSV format, No compressed option. From the custom columns, we choose 9 columns and arrange them as *Entry Name*, *Reviewed*, *Sequence*, *Pfam*, *Subcellular location [CC]*, *Gene Names*, *STRING*, *Organism*, and *Protein names*, as shown in **Figure S8**. The results reported for the filtered human/mouse dataset are generated from files of previous versions of STRING and UniProt, which are provided by the authors of the CFAGO paper, and those information are different from what we retrieve here for multi-species training.

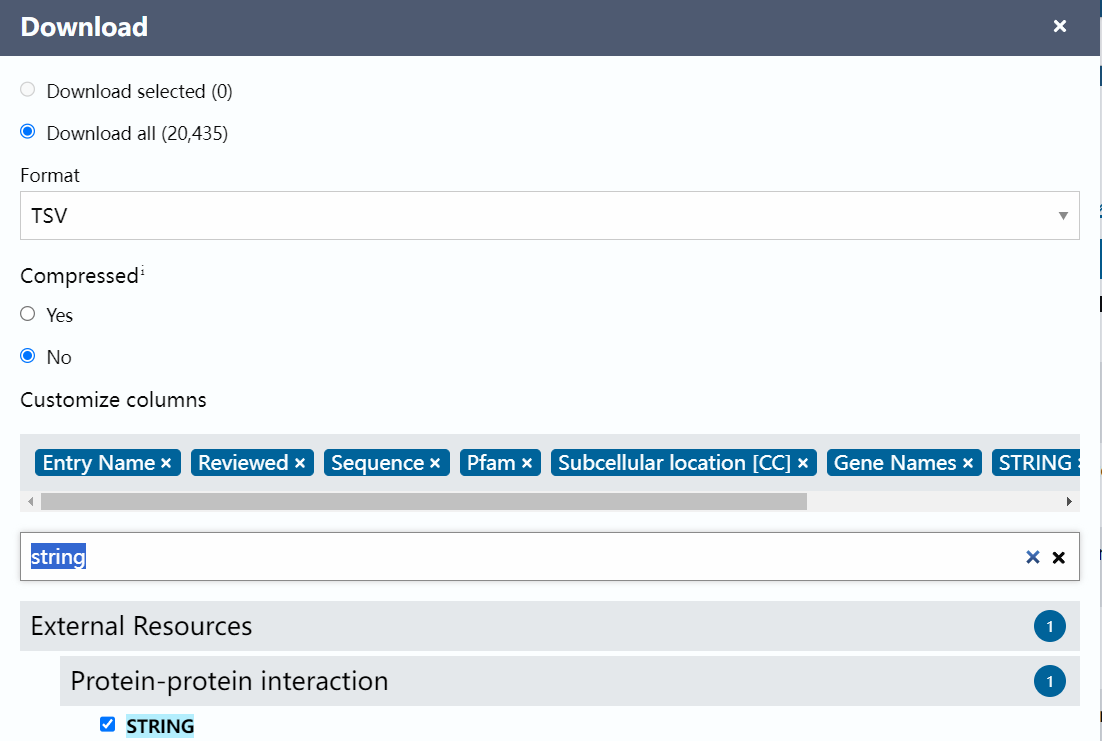

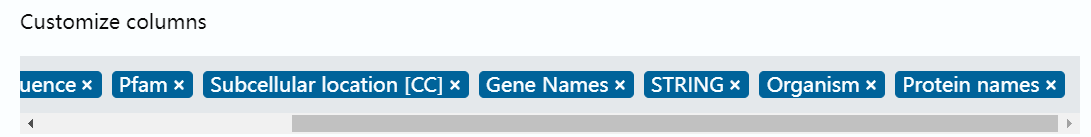

**Figure S8.** Download settings for UniProt annotations.

**C. Annotation for Gene Ontology**

Although we do not use GO annotation downloaded from the database but rather use those filtered and preprocessed by authors of the TEMPROT paper, we also provide a method to gather the GO annotation for those who are interested in training a DualNetGO for only one specific species. We recommend downloading the GO annotation from the QuickGO website. We first filter the database using a certain taxonomy code and only select those with one of the eight evidences described in the manuscript such as ‘IPI’, ‘EXP’. An example for downloading the filtered human GO annotations is provided in **Figure S9**.

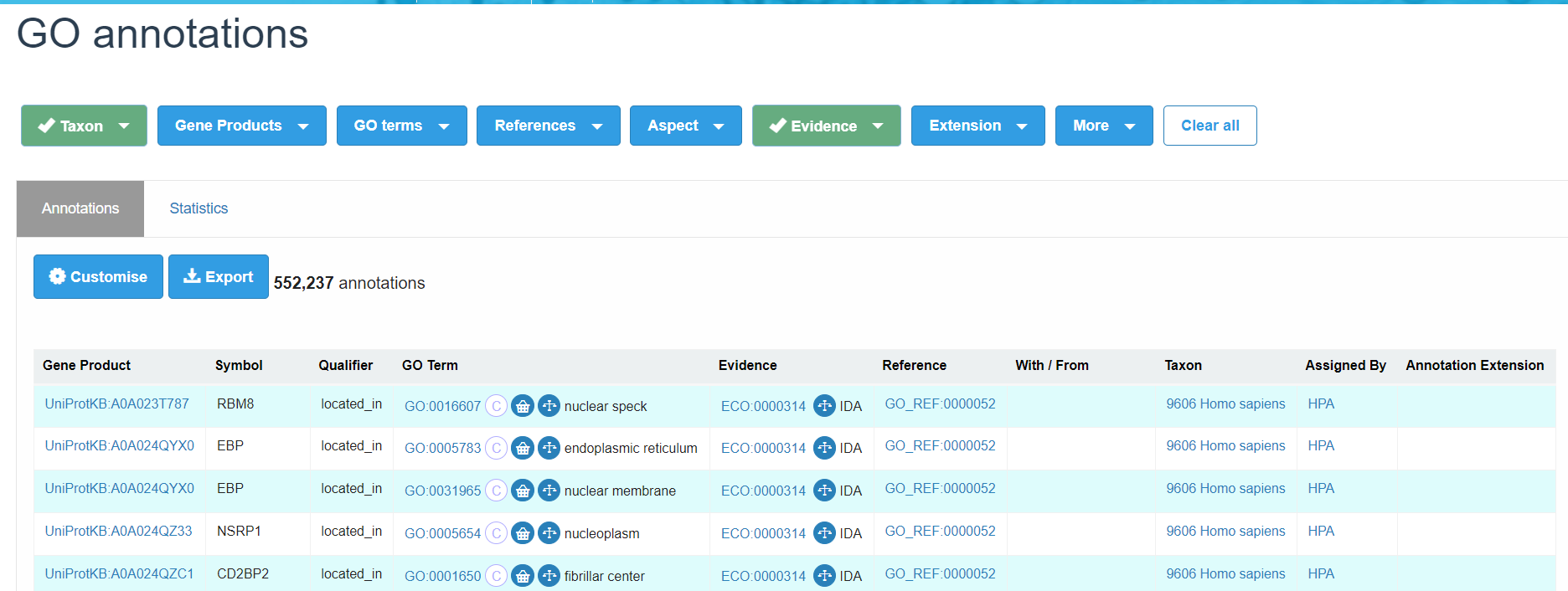

**Figure S9.** Download site of QuickGO for human GO annotations.

**16. Transformer Auto-Encoder training, graph features retrieval, and sequence embeddings retrieval**

For each species, a separate Transformer Auto-Encoder is trained and used to retrieve the hidden states for graph features. The data preprocessing and training procedures are mostly the same as those described in the manuscript. One difference is that for data in the latest version of STRING and UniProt, the original script used for the filtered human/mouse dataset (*attribute_data_preprocess.py*) is not compatible with the current format. We have provided a new script named *attribute_data_preprocess_new.py* to process the new data. Scripts for processing PPI networks and GO annotations remain unchanged. To save processing time, we set the number of training epochs for each network of each of the 15 species as 500 rather than 5000. Under this setting, one potential drawback is that 500 epochs may be insufficient to train the TransformerAE for graph-feature fusion, resulting in a worse quality of hidden states than those produced by a model trained for 5000 epochs. In **Table S12** we see that although most of the proteins in the PPI networks of human and mouse contain UniProt annotations, other species have much fewer annotations. As a result, the graph features from TransformerAE may be less informative given a sparse attribute matrix. After the self-supervised training of TransformerAE, there is an *npy* object saved in the output folder containing the hidden state matrix for all proteins in the PPI network of the specific species. Previously a protein STRING ID list for each species has been saved in the pickle format after the preprocessing of the PPI networks completes. The saved STRING IDs are consistent with those in the hidden state matrix. To further relate STRING IDs to external IDs such as UniProt entries, gene names, or protein names, one can use the protein information downloaded from the STRING database (*.protein.info.v12.0.txt.gz), use the UniProt annotation file, or use the UniProt ID mapping tools.

As we observe that, except for human and mouse, the UniProt Pfam and Subcellular location information do not cover a large portion of proteins in the PPI networks, these annotations may be unsuitable for general protein properties under the multi-species setting. Considering that we have discussed the potential to integrate multi-modal information in DualNetGO, we use a sequence embedding encoded by a protein language model Esm-2 as the final feature matrix in the feature selection space rather than the original Pfam/Subloc one-hot embedding matrix. As such, the hidden states from TransformerAE are still generated using the Pfam/Subloc attributes, but for the final feature selection we use the Esm2 embeddings instead. The embedding procedure is the same as that described in the TEMPROT paper. Weights of the Esm2 model can be directly downloaded from the *huggingfaces* website (**Figure S10**).

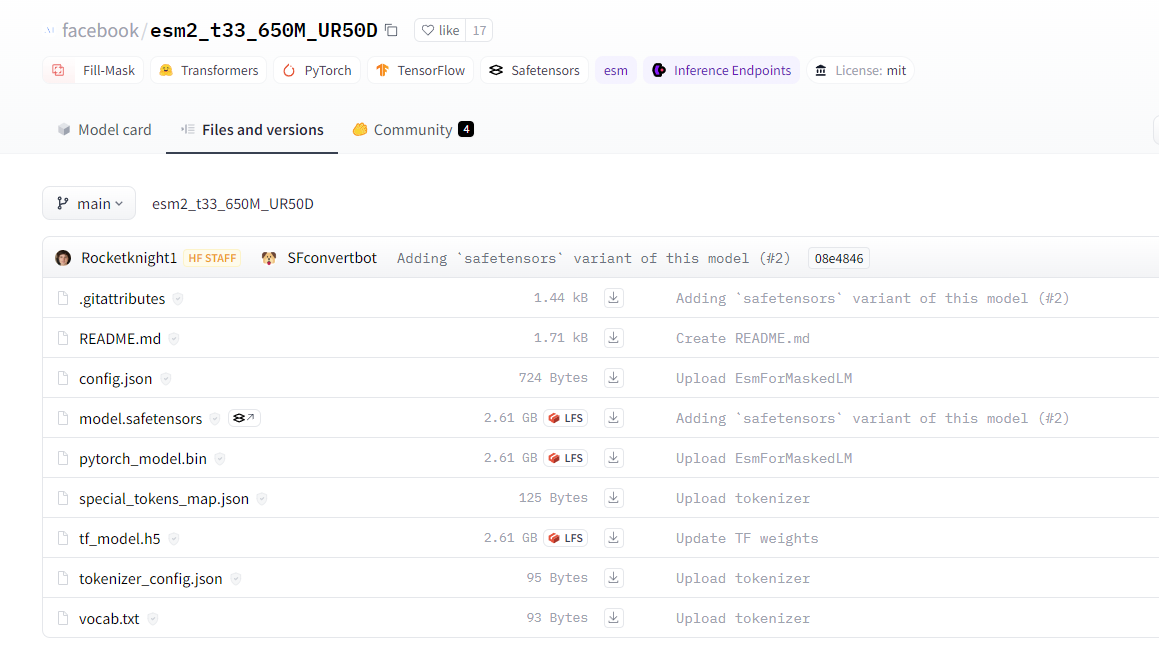

**Figure S10**. Download Esm2 model on *https://huggingface.co/facebook/esm2_t33_650M_UR50D/tree/main.*

For each sequence, a sliding window of 500 amino acids is used, and for any sequence longer than 500 aa, the sequence is split into chunks of length 500 with 250 aa as overlap. Every chunk is encoded separately and embeddings of all chunks are averaged to form the final embedding for a sequence. Sequence information can be retrieved with the *.protein.squences.v12.0.fa.gz file downloaded from the STRING database. The embedding matrix is arranged in the same protein order as the hidden state matrix. More details can be found on our github page.

**17. Training and evaluating DualNetGO with multi-species setting using CAFA3 training set and test set**

With hidden states of graph features and sequence features for all 15 species at hand, we can now train the multi-species DualNetGO model for the CAFA3 training set. The training, validation, and test set of CAFA3 are provided by the TEMPROT paper, and they are also used in other models such as DeepGOCNN, TALE, and Domain-PFP. Protein numbers in the CAFA3 test set are 2392, 1137, and 1265 for BP, MF and CC aspect, respectively, and the corresponding numbers of GO terms are 3992, 667, and 551.

Since not every protein in the CAFA3 training set has a record in the PPI networks, we only retain those included in the PPI network, resulting in a smaller training set than the original one. This step requires additional ID mapping from the UniProt Entry to STRING ID. The same procedure is applied to the validation set. For each protein in the training set, we first determine which of the 15 species it belongs to, then we do a SRTING ID mapping between the current protein and the protein ID list in the corresponding species-specific network and retrieve the specific-specific network index. After id mapping, each protein in the training set has a taxonomy code and a network index. Using these two pieces of information we can retrieve the corresponding hidden state embedding and the Esm2 sequence embedding for a protein. The same procedure is applied to each protein in the validation set. Finally, all graph hidden states and sequence embeddings are stacked together and totally eight feature matrices (seven network hidden state matrices and one sequence embedding matrix) are constructed as described for the filtered human/mouse dataset. With the corresponding one-hot encoded ground truth GO-term vectors as labels, we train the DualNetGO model with 0.001 learning rate for the Classifier and 0.01 for the Selector, using a dimension of 1024 for intermediate hidden layers in the Classifier. To ensure sufficient training, we set the number of epochs for Stage 3 of training DualNetGO as 1500, and apply an early stopping strategy with a patience epoch of 100. Other settings remain the same as for the filtered human/mouse dataset. For the testing set, we keep all proteins no matter if they are included in the PPI network, and perform BLASTp for all test proteins against all proteins we used to train TransformerAE models across 15 species. Only the target protein with the highest identity is retained for each query protein in the test set, and the E-value threshold is set as 10 so that we ensure a 100% query rate. The corresponding feature matrices of target proteins are used as those for query proteins so that our model can predict functions for any proteins even for those not seen by the PPI network. Fmax and AUPR are used as metrics to evaluate the prediction performance of our model.

In addition, to utilize homology information, we integrate the homology search procedure using BLASTp. Specifically, we perform BLASTp for proteins in the CAFA3 test set against the unfiltered CAFA3 training set (aspect-specific) and attain the identity scores. The GO-term labels of target proteins are used as the labels of query test proteins, with identity scores as coefficients. To further balance the homology score and the predicted score, we multiply the homology scores with the maximum value of the predicted scores. Two coefficients are designed based on the results on the validation set to combine the homology score and DualNetGO predicted score, and the two coefficients sum up to 1. We name the coefficient for DualNetGO score as **alpha** and the one for homology score as **1-alpha**.

$$Score_{ensemble} = alpha\times Score_{DualNetGO}+max(Score_{DualNetGO})\times(1-alpha)\times Score_{BLASTp}$$

To distinguish between the DualNetGO model with and without homology search, we name the one with homology search as **DualNetGO**+ and the other **DualNetGO**.

Hyperparameters for reproducing the results reported in the paper are shown in **Table S14**.

| Training set | Aspect | E1 | E2 | E3 | $N_{f}$ | Alpha |
| --- | --- | --- | --- | --- | --- | --- |
| CAFA3 | BP | 500 | 10 | 1500 | 2 | 0.72 |
|  | MF | 300 | 70 | 1500 | 2 | 0.58 |
|  | CC | 400 | 10 | 1500 | 2 | 0.82 |

**Table S14.** Hyperparameter settings of DualNetGO for CAFA3 dataset.

Codes for the training and testing procedure are provided at the DualNetGO github site.

**18. Evaluation of DualNetGO trained on CAFA3 data on the filtered human/mouse dataset**

We also evaluate the performance of other state-of-the-art models on the filtered human/mouse dataset. These models utilize various information sources such as sequences and structures, and they are trained on the CAFA3 or SwissProt multi-species datasets.

**Table S15** shows that DualNetGO that trained on the filtered dataset (the 5^th^ row) performs the best on the CC aspect, produces the second best results on BP and CC, and gives worse results on the MF aspect than some others.

However, the comparison is somewhat unfair. As single-species DualNetGO models are only trained on single-species datasets, they encounter significantly fewer sequences than other models during the training process, which is a great disadvantage compared with those utilize the homology search strategy such as NetGO3.0 and DeepGOplus. In addition, we use the pretrained model provided by DeepGOplus and the online server of NetGO3.0 to perform the evaluation but do not re-train them only using the filtered human/mouse dataset, making it hard to do a fair comparison. Even though DualNetGO is trained on significantly fewer GO terms whereas others are trained on hundreds and thousands of GO terms, the filtered GO terms are mostly included in the CAFA3 dataset. Training with more sequences not only allows the model to explicitly adopt homology search but also implicitly encode information from similar sequences. However, when we test our multi-species DualNetGO model, which is trained on the CAFA3 data, on the single-species human/mouse datasets, we still get worse results on the BP and MF aspect than NetGO3.0 and DeepGoplus. One possible reason is that the MF and BP aspects are more related to sequence properties which are mainly exploited by NetGO3.0 and DeepGOplus, whereas CC is more related to PPI network properties, as discussed by another study entitled “Protein function prediction as approximate semantic entailment” by Kulmanov et. al. published in 2024.

The good results produced by the multi-species network-based model DeepGraphGO indicate that, by end-to-end training a model utilizing network information from multi-species via GNN models, it is feasible to produce comparable results on the BP and MF aspects. Our DualNetGO model currently cannot be trained in an end-to-end manner, so the prediction accuracy completely depends on the overall qualities of the hidden states from the self-supervised learning of TransformerAE. However, both the network adjacency matrix and the Pfam/Subloc attribute matrix could be very sparse for some species, which could result in less informative graph embeddings. From another aspect, the non-end-to-end training procedure makes the DualNetGO model a flexible one to include information from other sources and make the best out of all combinations of these features.

In summary, the DualNetGO model produces the best results on the CC aspect even when trained with much fewer sequences than other models, and also outperforms other multi-species models when trained with the CAFA3 dataset. The final integrated graph embeddings produced by DualNetGO could be further used as features to improve the performance of other ensemble models such as NetGO3.0.

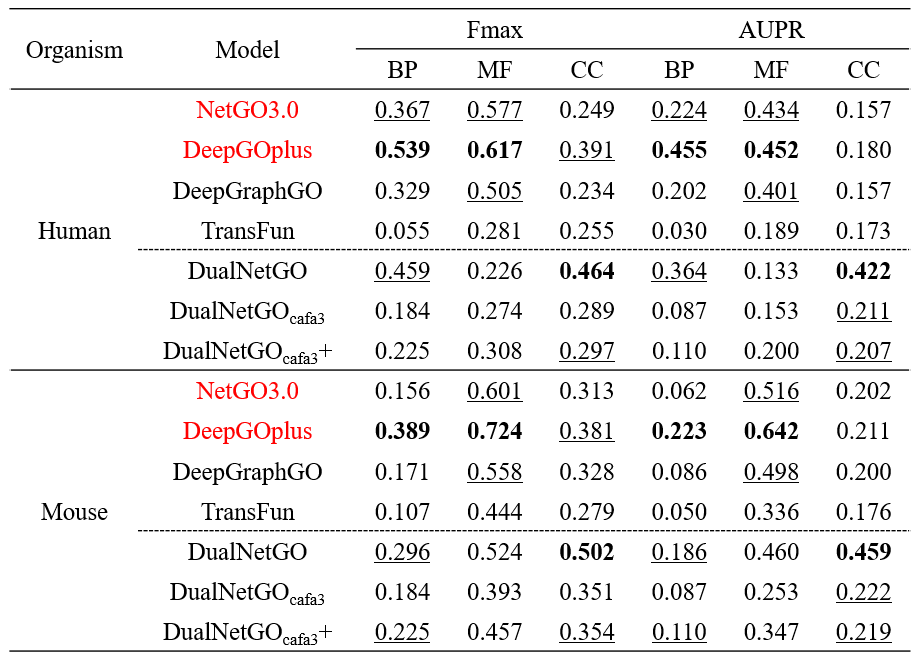

**Table S15.** Comparison of DualNetGO with other SOTA methods on protein function prediction using the filtered human/mouse dataset as the test set. Methods that utilize the homology search strategy are marked by red color. DualNetGO refers to the model trained on the filtered single-species dataset; DualNetGO_cafa3_ and DualNetGO_cafa3_+ refer to models trained on the CAFA3 multi-species dataset without and with homology search, respectively. Best results are marked in bold, and second and third best results are underlined.

**19. Feature selection results of DualNetGO on human/mouse and CAFA3 datasets**

As DualNetGO can determine a suitable subset of PPI for protein function prediction, the chosen combination reflects the importance of different PPI networks.

**A. On filtered human/mouse datasets**

For the embedding-centric model, we implement five different graph embedding algorithms and count the occurrences of each PPI network in these experiments. As shown in **Figure S11a**, *textmining* network provides the most valuable information in BP prediction for both human and mouse, and CC is more related to protein attributes (denoted as feature) than other features. CC's close relation to protein attributes is not surprising because the protein attributes used in this study include Pfam domain and subcellular location, and subcellular location could cause information leakage due to its close relation to CC. In the evidence-centric model, the number of graph embeddings chosen as features is counted across different PPI networks. In **Figure S11b** we observe that TransformerAE generally performs better than the other methods, demonstrating its superior performance. However, the simpler deep learning autoencoder, MLPAE, is also effective in extracting PPI information for MF prediction.

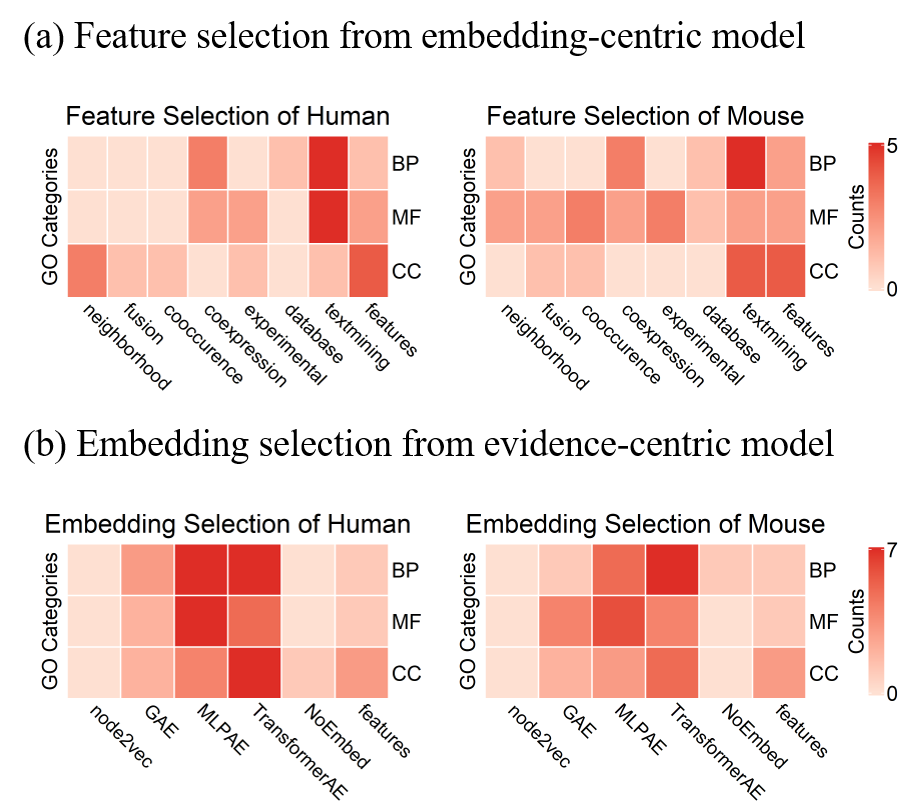

**Figure S11**. Feature selection across different settings of DualNetGO in the filtered human/mouse dataset.

**B. On the CAFA3 dataset**

We perform a thorough hyperparameter search on E1, E2 and num_feat_select on all three aspects, and retrieve the features selected by DualNetGO. We count the frequency of each feature matrix being selected by DualNetGO models with top-10 Fmax scores for BP, MF and CC aspect.

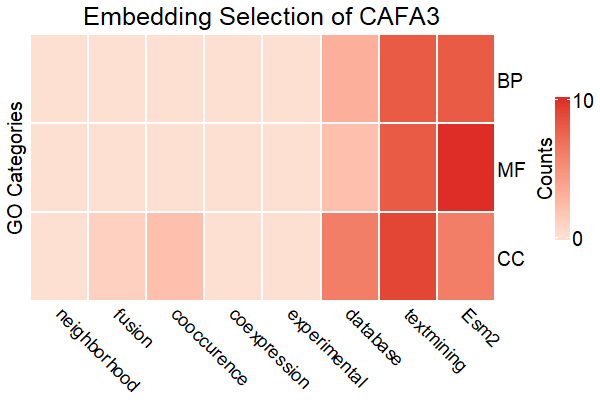

**Figure S12**. Feature selection across different hyperparameters of DualNetGO in the CAFA3 dataset.

Results are shown in **Figure S12**. For BP the best result is produced by combining the hidden states from the *textmining* network and Esm embeddings. For MF the best result is produced by combining the hidden states from the *database* network and Esm embeddings. For CC the best result is produced by combining the hidden states from *cooccurrence* and *textmining* networks. This phenomenon also suggest that CC is more related to the PPI network property than the sequence property.

**20. Effects of the “selection during training” strategy**

The DualNetGO model is trained by alternate evaluation of feature selection and update of classifier’s weights. Before the best feature combination is determined, all features contribute to the classification loss during the feature sampling process. Therefore, even though a certain feature matrix is not included in the final features and not used for prediction, it helps update the classifier’s weights in stage 1 and stage 2, and attributes to the improvement of the model. This gives DualNetGO the advantage of fully utilizing all collected data for training the classifier but only using an optimal subset of them for prediction. To demonstrate the effectiveness of the “selection during training” strategy, we enumerate all possible combinations of feature matrices and train a corresponding classifier using each of the combinations. Results in **Table S16** show that the best results from all feature combinations (denoted as **Enumerate**) are worse than those generated from DualNetGO (denoted as **Select**) in the filtered human/mouse datasets. Similar results are found in the CAFA3 multi-species dataset except a slightly higher AUPR socre on the MF aspect for the Enumerate model, shown in **Table S17**.

| Feature | Model | Human | | | Mouse | | |
| --- | --- | --- | --- | --- | --- | --- | --- |
|  |  | BP | MF | CC | BP | MF | CC |
| UniProt | Select | 0.459 | 0.226 | 0.464 | 0.296 | 0.524 | 0.502 |
|  | Enumerate | 0.334 | 0.199 | 0.379 | 0.262 | 0.478 | 0.490 |
| Esm2 | Select | 0.455 | 0.232 | 0.487 | 0.304 | 0.545 | 0.506 |
|  | Enumerate | 0.306 | 0.194 | 0.346 | 0.253 | 0.468 | 0.434 |

**Table S16.** Comparison between Fmax scores from DualNetGO feature selection training and from the best results out of all feature combinations on the filtered datasets.

| Model | Fmax | | | AUPR | | |
| --- | --- | --- | --- | --- | --- | --- |
|  | BP | MF | CC | BP | MF | CC |
| Select | 0.565 | 0.601 | 0.691 | 0.576 | 0.619 | 0.738 |
| Enumerate | 0.535 | 0.591 | 0.689 | 0.531 | 0.620 | 0.721 |

**Table S17.** Comparison between performance from DualNetGO feature selection training and from the best results of all feature combinations on the CAFA3 test set.

Noted that the optimal combination of features selected by the “selection during training” strategy of DualNetGO is not the same as the set of features that are found by enumeration to provide the best results. Most of them are overlapped, indicating that the DualNetGO model plays the feature selection role (**Table S18, 19**). Comparing the features selected by DualNetGO and by Enumerate when using Esm2 embeddings as protein attributes instead of the original Pfam+subloc information retrieved from UniProt, we find that the same set of features are selected for the CC aspect on the human dataset. However, the performance of DualNetGO is better than that of Enumerate, which directly demonstrates the importance of incorporating other features in the training process for better performance. In addition, we find that using Esm2 embeddings instead produces higher results than using UniProt annotations by DualNetGO except on the BP aspect of human, but Esm2 is only selected as the final feature for human CC and mouse MF. Furthermore, the Esm2 embeddings themselves may not be as predictive as the original UniProt annotations, as the best results from enumerating combinations of Esm2 embeddings are worse than those from enumerating combinations of UniProt annotations. These phenomena suggest that features of lower relevance can also be utilized by DualNetGO to improve its performance.

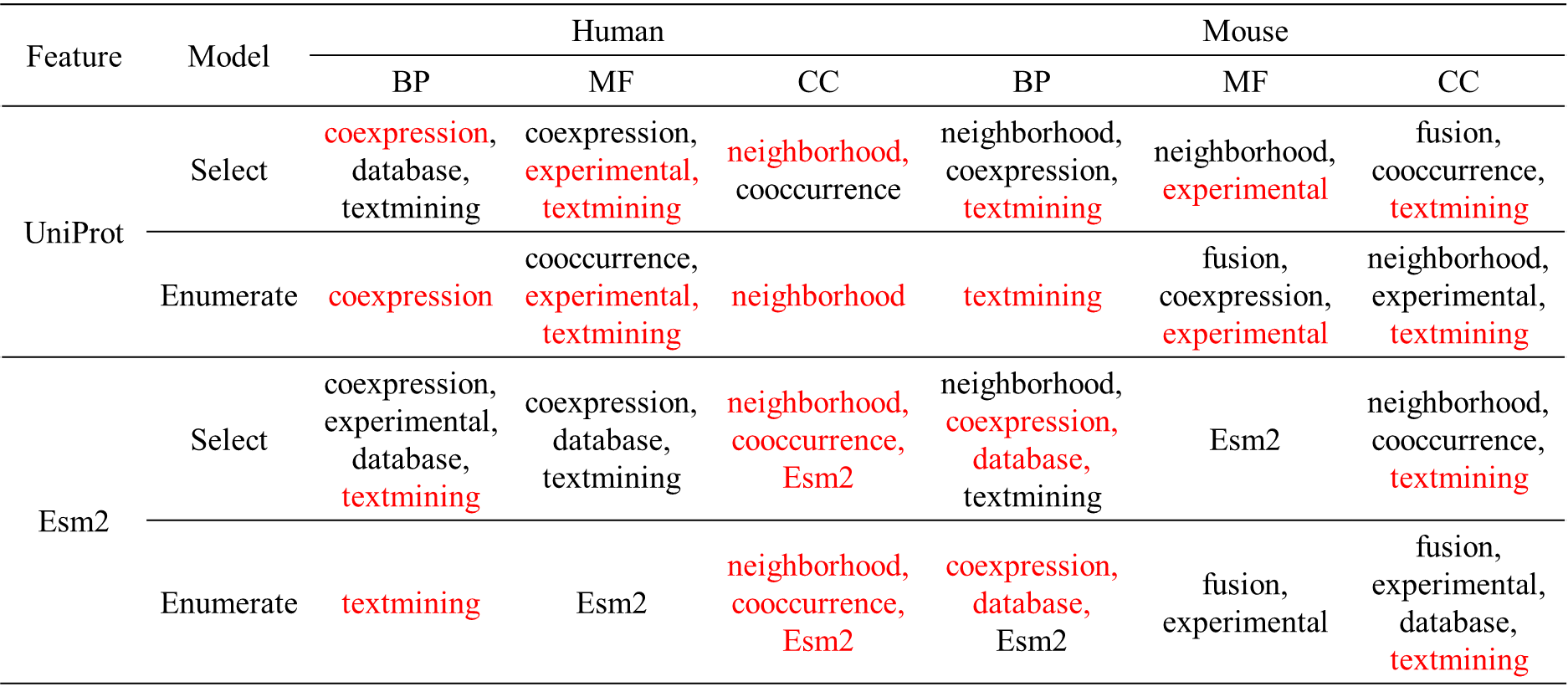

**Table S18**. Selected features by DualNetGO (Select) and by choosing from the best results across all combinations (Enumerate) of the filtered human/mouse datasets. Overlapping features are in red.

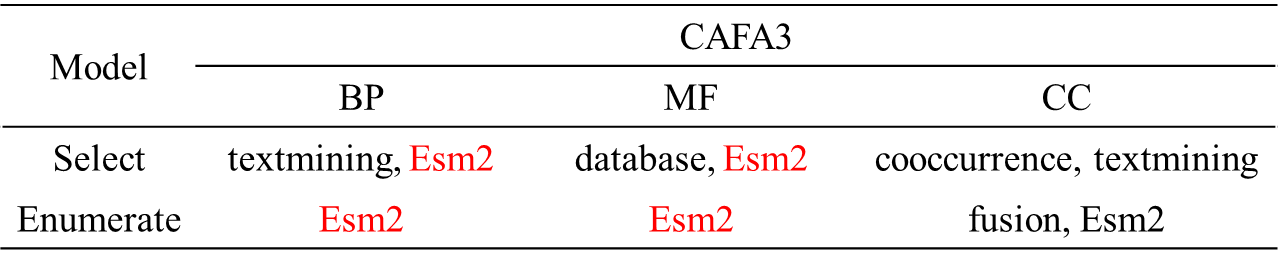

**Table S19**. Selected features by DualNetGO (Select) and by choosing from the best results across all combinations (Enumerate) of the CAFA3 test set. Overlapping features are in red.
